# Engineered yeast multicellularity via synthetic cell-cell adhesion and direct-contact signalling

**DOI:** 10.1101/2024.06.23.600301

**Authors:** Fankang Meng, William M. Shaw, Yui Kei Keith Kam, Tom Ellis

## Abstract

Coordination of behaviour in multicellular systems is one the main ways that nature increases the complexity of biological function in organisms and communities. While *Saccharomyces cerevisiae* yeast is used extensively in research and biotechnology, it is a unicellular organism capable of only limited multicellular states. Here we expand the possibilities for engineering multicellular behaviours in yeast by developing modular toolkits for two key mechanisms seen in multicellularity, contact-dependent signalling and specific cell-to-cell adhesion. MARS (<u>M</u>ating-peptide <u>A</u>nchored <u>R</u>esponse <u>S</u>ystem) is a toolkit based on surface-displayed fungal mating peptides and G protein-coupled receptor (GPCR) signalling which can mimic juxtacrine signalling between yeasts. SATURN (<u>S</u>accharomyces <u>A</u>dhesion <u>T</u>oolkit for multicell<u>U</u>lar patte<u>RN</u>ing) surface displays adhesion-proteins pairs on yeasts and facilitates the creation of cell aggregation patterns. Together they can be used to create multicellular logic circuits, equivalent to developmental programs that lead to cell differentiation based on the local population. Using MARS and SATURN, we further developed JUPITER (<u>JU</u>xtacrine sensor for <u>P</u>rotein-protein In<u>TER</u>action), a genetic sensor for assaying protein-protein interactions in culture, demonstrating this as a tool to select for high affinity binders among a population of mutated nanobodies. Collectively, MARS, SATURN, and JUPITER present valuable tools that facilitate the engineering of complex multicellularity with yeast and expand the scope of its biotechnological applications.

## Introduction

The transition from unicellular to multicellular organisms represents a pivotal point in biological evolution, driven predominantly by three traits: intercellular communication, cell adhesion, and cell differentiation^1^. Together these mechanisms enable populations of cells to coordinate behaviour and achieve more complex functions than single-celled systems. *Saccharomyces cerevisiae* yeast is one of the most widely-used organisms in research and industrial biotechnology, but has a largely unicellular lifestyle with no native capacity for multicellular complexity. Modular DNA toolkits exist for rapid, advanced genetic engineering in yeast that can provide the equivalent of differentiation, where cells given external inputs change their gene expression to take on new specialist tasks^2–5^. However, no equivalent toolkits exist for programmable cell-to-cell adhesion in yeast and intercellular communication tools have so far focused only on those with diffusible signals, such as plant cytokinin^6^ and animal hormones^7^.

In multicellular systems, intercellular communication is key to a wide range of cellular behaviors^8^. It is primarily divided into four categories: autocrine, paracrine, endocrine, and juxtacrine signalling. The first three involve the secretion and diffusion of signalling molecules, and multiple genetic tools have been developed to endow engineered cells to be programmed do synthetic cell-cell signalling of this kind^9–12^. Juxtacrine signalling, however, is the most relevant kind for multicellular organisms and requires direct cell-cell contact where cell surface ligands, such as peptides or proteins, engage with receptor proteins on adjacent cells via proximity, without the signals diffusing away from the source cell. For example, the synNotch system, developed for use in mammalian cells, is activated by cell-cell direct interactions, which in turn prompts cells to express adhesion proteins and aggregate into specialized multicellular structures^13,14^.

Although engineered cell contact-based signalling has been widely demonstrated in mammalian cell systems, it has not been exploited for use in yeast, despite yeast being important for high-throughput research in investigating biological mechanisms and for drug development^15,16^. In past work, yeast surface display technologies have been used to establish cell contact-based signalling associations directly between yeast and mammalian cells to explore immune-related protein interactions^17,18^. However, a juxtacrine-like signalling system that works exclusively between yeast cells has not been described and could offer significant value for increasing the complexity of synthetic biology, especially if used in engineered populations of yeast cells that can have been designed to specifically bind to one another.

GPCR signalling is extensively used in eukaryotes as a method for cells to sense and respond to their extracellular environment^19^. GPCRs sense a diverse range of signals by undergoing conformational changes upon ligand binding^20^ and in fungi like yeast, GPCRs regulate highly specialized mating behaviours via the sensing and binding of signal peptides, like the α-factor of *S. cerevisiae*^21^. The α-factor is a short peptide secreted by yeast and recognized by the Ste2 GPCR protein in the cell membrane of other yeast cells (**Fig. 1a**). Once recognised by the GPCR, this then triggers an intracellular signalling cascade in the yeast that activates gene expression changes. GPCRs derived from mammalian and fungal sources have been successfully utilised in yeast to regulate cell-to-cell communication and behaviour through signal secretion^9^. Here we anticipate that GPCR signalling could also provide a framework for juxtacrine signalling if we can engineer the availability of signalling peptide ligands to be dependent on direct cell-cell contact. Combining this with a controlled system for synthetic and orthogonal cell-to-cell adhesion that directs yeast cells in a population to aggregate in defined patterns offers a route to multicellularity.

**Fig. 1.**
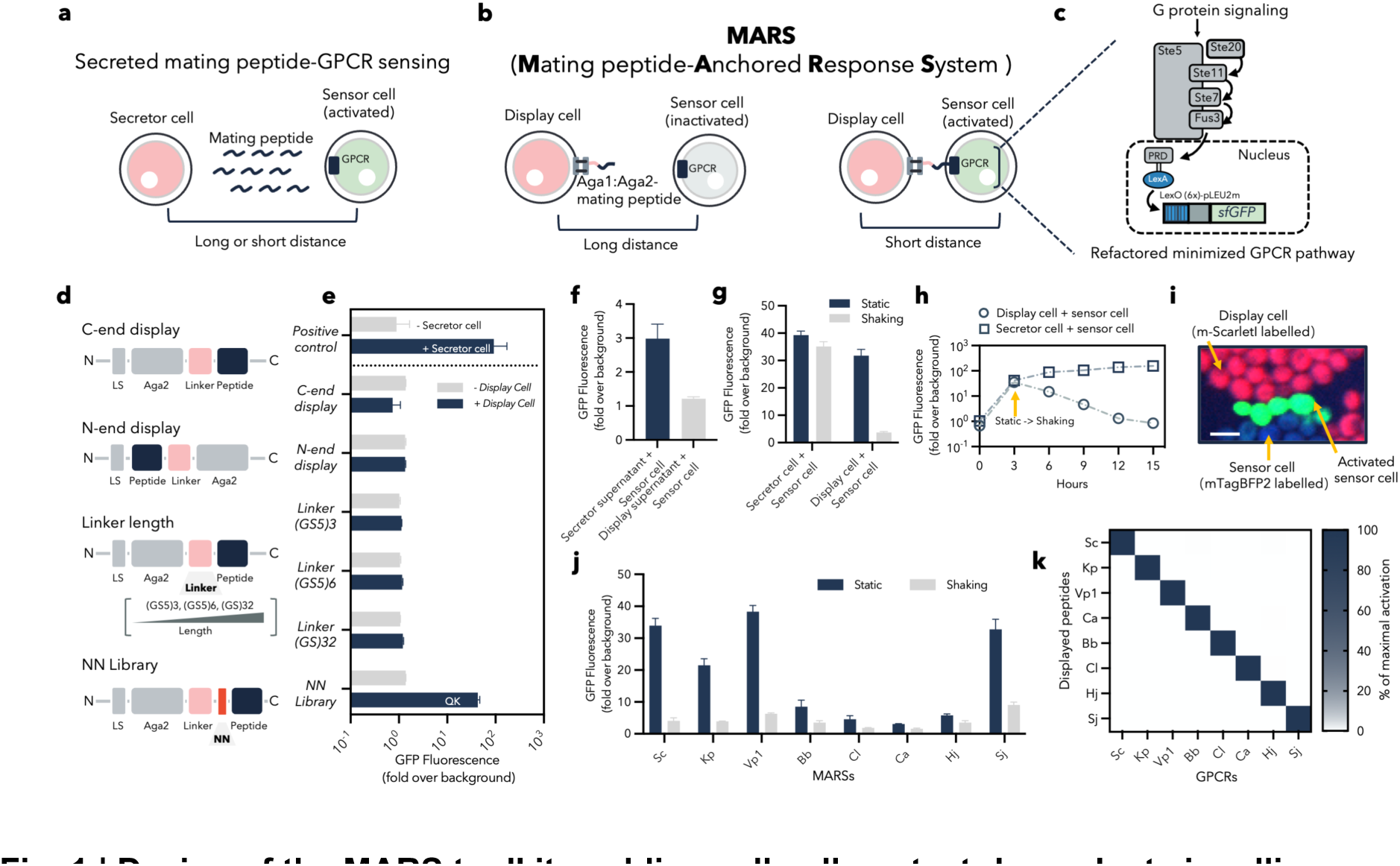
| Design of the MARS toolkit enabling cell-cell contact-dependent signalling. **a** Schematic representation of secreted mating peptide-GPCR sensing. Secreted peptide from the secretor cell can activate the sensor cell across both short and long distances. **b** Schematic representation of the MARS design utilising Aga1-Aga2 yeast surface display system. Interaction of a displayed peptide from the display cell with its specific GPCR (Ste2) on the cell surface initiates gene activation in the sensor cell. **c** Upon recognition of the displayed ligand by the membrane-bound Ste2 GPCR, the signal then goes through the MAP-kinase-mediated phosphorylation cascade to ultimately activate a synthetic promoter, LexO(6x)-pLEU2m, via the LexA-PRD synthetic transcription factor, generating a green-fluorescent signal as a measurable readout. **d,e** Schematic (**d**) and characterisation (**e**) of four strategies to display Sc_peptide on the yeast cell surface; data was recorded at the 4 hour time point. LS = leader sequence. **f** Characterisation of the activation level of sensor cells after co-culturing with overnight culture supernatant from secretor and display cells; data were recorded at the 4-hour time point. **g** Characterisation of the activation level of the sensor cell after co-culturing with secretor and display cells under shaking (700 RPM) and static conditions; data were recorded at the 4-hour time point. **h** Characterisation of the activation level of the sensor cell over time after co-culturing with secretor and display cells. Initially cultured under static conditions, the system was then shifted to shaking conditions (700 RPM). The OD600 of the culture, post-adjustment to shaking conditions, was kept equivalent to that at the 3-hour time point by dilution at each data acquisition time point. **i** An example microscopy image illustrating cell-cell contact-dependent activation of the sensor cell in the Sc_MARS system. **j** Characterisation of the activation level of 8 different MARS systems under shaking (700 RPM) and static conditions; data recorded at the 4-hour time point. **k** Characterization of orthogonality among all 8 MARSs; data recorded at the 4-hour time point under static conditions in a 96 deep-well plate. All experiments were conducted in triplicate, and error bars represent standard deviation.

In this work, we enable programmable multicellular behaviours in yeast by developing two new genetically encoded toolkits for *S. cerevisiae*. We first establish juxtacrine-like signalling that is dependent on cell-to-cell contact using a toolkit called MARS (Mating peptide-Anchored Response System), which leverages yeast surface display to anchor different signalling peptide ligands to yeast’s cell surface with these recognised by GPCRs expressed by other yeast cells. A set of 8 highly orthogonal pairs were developed for this. We then created a highly-engineered yeast strain optimised for introducing synthetic adhesion and for this developed a synthetic cell-to-cell binding toolkit, termed SATURN (Saccharomyces Adhesion Toolkit for mUlticellular patteRNing) that can direct cells in a population to bind selectively to one another with different affinities. We combined SATURN with MARS to achieve multicellular behaviours akin to development, and complex logic, before finally demonstrating the design of a novel genetic sensor for protein-protein interactions termed JUPITER (JUxtacrine sensor for Protein-protein InTERaction).

## Results

### Establishing cell-to-cell contact-based signalling

To achieve the design of a contact-based GPCR signalling system, we reasoned that by anchoring signal peptides via yeast cell surface display, the communication process between the displayed signal peptides and Ste2 GPCR proteins will depend on direct cell-cell contact. For this we established the MARS system using the mating pheromone α-factor peptide of *S. cerevisiae* (Sc_MARS). Sc_MARS contains two types of engineered yeast cells, namely display cells and sensor cells (**Fig. 1b**). We designed both cells to be based on a prior yeast strain developed by us with a refactored GPCR-signalling pathway from tunable sensing^7^ (**Fig. 1c**) and deleted the native *aga1* and *aga2* genes in this strain via CRISPR-Cas9 to create yWS2064 (**Tables S1**). For the display cell, we first used the Aga1-Aga2 yeast cell surface display technique (C-end display) to attach the α-factor peptide (Sc_peptide) to the yeast surface. To link the Aga2 domain to the Sc_peptide in a fusion protein, we first tried a flexible linker consisting of eight glycine-serine repeats, (GS)8. For the sensor cell, the *S. cerevisiae* Ste2 GPCR (Sc_GPCR) was expressed from a constitutive promoter with its activation linked to a fluorescence output. Once membrane-bound Sc_GPCR recognizes Sc_peptide, a MAP-kinase-mediated phosphorylation cascade occurs in the sensor cell that activates the synthetic promoter, LexO(6x)-pLEU2m, via the LexA-PRD synthetic transcription factor, and triggers GFP expression.

We first tested functionality of this initial Sc_MARS design in a standardized workflow. We simulated cell-cell contact by co-culturing sensor and display cells together under two conditions: a static condition where cells can have close contact with each other; and a shaking condition where cells cannot maintain physical contact. If Displayed peptide-receptor signalling is truly cell contact dependent, an obvious difference should be seen in sensor cell activation levels between these conditions that is not seen when the peptide is secreted (**Fig. S1**) rather than displayed.

Initial results failed to show an expected increase in sensor cell activation in static co-culture experiments with cells displaying peptide, presumably due to ineffective design in terms of how it is displayed on the yeast cell wall and whether that restricts its ability to contact the Ste2 GPCR on a different cell. To address this, various strategies were employed to optimise peptide display (**Fig. 1d**). Initially, alterations were made to the peptide’s display orientation on the Aga2 protein, switching from a C-terminal to an N-terminal presentation for the Sc_peptide. However, neither version activated sensor cells. Subsequently, we varied the linker length between Aga2 and the signal peptide region by testing flexible glycine-serine peptide linkers of varying lengths: (GS5)_3_, (GS5)_6_, and (GS)_32_. However, no increase sensor cell activation was seen. Lastly, we focused on the amino acid composition immediately preceding the peptide in the Aga2-peptide fusion, hypothesizing that it could affect peptide structure or its ability to interact with Ste2 GPCR^22^. Employing what we termed the ’NN strategy,’ we randomly altered the two preceding amino acids and screened for combinations that could activate the sensor cells. Via DNA assembly with degenerate primers, we constructed a variant library with general structure Aga2-N-N-Sc_peptide (where N = any amino acid). We then characterised variants by randomly selecting colonies and co-culturing with sensor cells (**Fig. S2a**). This led to the identification of a variant with QK preceding the Sc_peptide that successfully activated Sc_GPCR sensor cells (**Fig. 1e** and **Fig. S2b**).

Given that Q and K are both charged amino acids, we first needed to rule-out that activation was just due to Sc_peptide being cleaved away from the Aga2 domain by native yeast proteases, leading to its secretion. Literature research and database reviews^23,24^ first gave us confidence that a QK-containing design would not contain high efficiency sites for native *S. cerevisiae* proteases (**Table S8**). However, despite this, the possibility remained that interplay of proteases in yeast’s secretory pathway could still result in peptide release. To experimentally test this, we collected overnight culture supernatants from the display cells and exposed these to the sensor cells. Notably this did not measurably activate sensor cells, contrasting with a control where supernatants containing peptides from secreting cells gave significant sensor cell activation (**Fig. 1f**). Further analysis under different co-culture conditions revealed that when display cells and sensor cells were co-cultured statically there was substantial activation of sensor cells and this activation showed a 8.6-fold decreased when in shaking conditions that inhibit cell-cell contact. Only a slight difference was seen between shaking and static condition when secretor cells were used instead (**Fig. 1g**). In further test, cell co-cultures were first started in static conditions and then switched to a shaking condition, and this resulted in a clear distinction of sensor activation level between the two co-cultures. In the co-culture with display cells, sensor cell activation decreased gradually, while in the co-culture with secretor cells, sensor cell activation continued to increase (**Fig. 1h**). Microscopy also confirmed that the interaction necessary for activation between sensor and display cells was dependent on direct cell-cell contact (**Fig. 1i**). Together, these findings affirmed that sensor cell activation from the optimised surface-displayed Sc_peptide is a result of cell-cell contact-dependent signalling, occurring exclusively when sensor and display cells achieve direct physical contact.

### Expanding the MARS toolkit for orthogonal communication

The development of diverse orthogonal cell communication tools is critical for achieving more complex systems in synthetic biology. In pursuit of this aim, we next expanded MARS by evaluating the performance of eleven additional GPCR sensors shown previously to function in *S. cerevisiae*^9^. This involved replacing the displayed peptide in the display cell and the Ste2 GPCR in the sensor cell with different peptide peptide-GPCR pairs. To optimise peptide display, we explored the same four strategies for each pair as we had done so for the Sc_peptide (**Fig. S2** and **Fig. S3**). Notably, the NN strategy yielded the highest success rates, with the QK motif succeeding for several MARSs, and suitable NN motifs found for others (**Fig. S2c-f**). Pairs not showing any effective activation after these strategies were abandoned. Following peptide-display optimisation for 7 new pairs (**Table S9**), we could demonstrate contact-dependent signalling for a total of 8 MARS pairs with signalling leading to a wide range of sensor cell activation profiles (**Fig. S4**).

We further evaluated the contact dependence of MARS to examine how the new peptide-receptor pairs behaved under different physical conditions. We found that Vp1 and Kp_MARS showed high activation in shaking in these tests. However, no detectable activation was seen when transferring supernatant from display cell culture to sensor cells, suggesting this was not due to peptide release (**Fig. S5a, b**). This led us to hypothesize that the observed effects for Vp1 and Kp may instead be due to an increased sensitivity caused by overexpression of the displayed peptide or GPCR^9^. To address this, we varied the expression levels of the displayed peptides and GPCRs by exchanging the promoters for their genes to ones of varying strengths and then selected the promoter combinations that led to higher dynamic range between static and shaking condition (**Fig. S5c**). For the best promoter combination in the Vp1 MARS optimisation, the dynamic range between static and shaking conditions increased from 1.1-fold to 6.1-fold, and sensor cell fluorescence level under shaking conditions (*i.e.* background) was successfully decreased by 17-fold. (**Fig. S5d, e**).

The same approach to tune peptide and GPCR expression levels was applied for Kp_MARS. Initial results showed a modest improvement in dynamic range, from 1.1-fold to 1.9-fold activation, and a 5.5-fold reduction in shaking cell fluorescence, however, this was not deemed sufficient. So, to further optimise the Kp_MARS we reintroduced the Sst2 negative feedback protein, a GTPase-activating protein previously deleted in our background strain due to its known role in downregulating the sensitivity of the mating-related GPCR pathway^7^ (**Fig. S5f**). This modification considerably improved the dynamic range to 5.5-fold activation in static compared to shaking and decreased the fluorescence under shaking by 21-fold (**Fig. S5g**). Furthermore, an obvious cell-cell contact dependent activation process was also observed by microscopy (**Fig. S5h**). We anticipate that these optimisation strategies can also be applied to other MARS pairs, for example to help avoid rapid activation due to transient cell-cell contacts. This could improve the precision and reliability of MARSs in complex synthetic biology applications.

After optimisation, all MARS peptide-receptor pairs demonstrated clear activation differences between static and shaking conditions (**Fig. 1j**). Importantly all eight pairs exhibited a significant level of orthogonality when cross-tested against each other (**Fig. 1k**).

### Design of multicellular logic based on MARS

Single organisms frequently encounter engineering constraints due to their limited capacity for exogenous DNA and the burden of running complex genetic circuits^25,26^. Multicellular microbial consortia, designed to cooperate and distribute tasks, offer a promising avenue for overcoming these limitations and engineering complex behaviours^27,28^. To showcase the opportunities to use MARS for this, we demonstrated several multi-cell logic circuits. We first designed a scalable chain topology that links signals between cells in a defined sequence. Each chain is initiated by a display cell that activates a sensor cell to then express and display a new ligand. This in turn activates another sensor cell to display a new ligand, and so on until the chain terminates with a final sensor cell (**Fig. 2a**). Experimentally, we showed that this scalable chain topology allowed for sequential signal transmission within a multicellular community up to six members (**Fig 2b, c**). We observed that subjecting a cell chain to shaking conditions or to the removal of a cell link disrupted signal continuity, demonstrating the chain’s reliance on contact-based signalling for signal propagation.

**Fig. 2.**
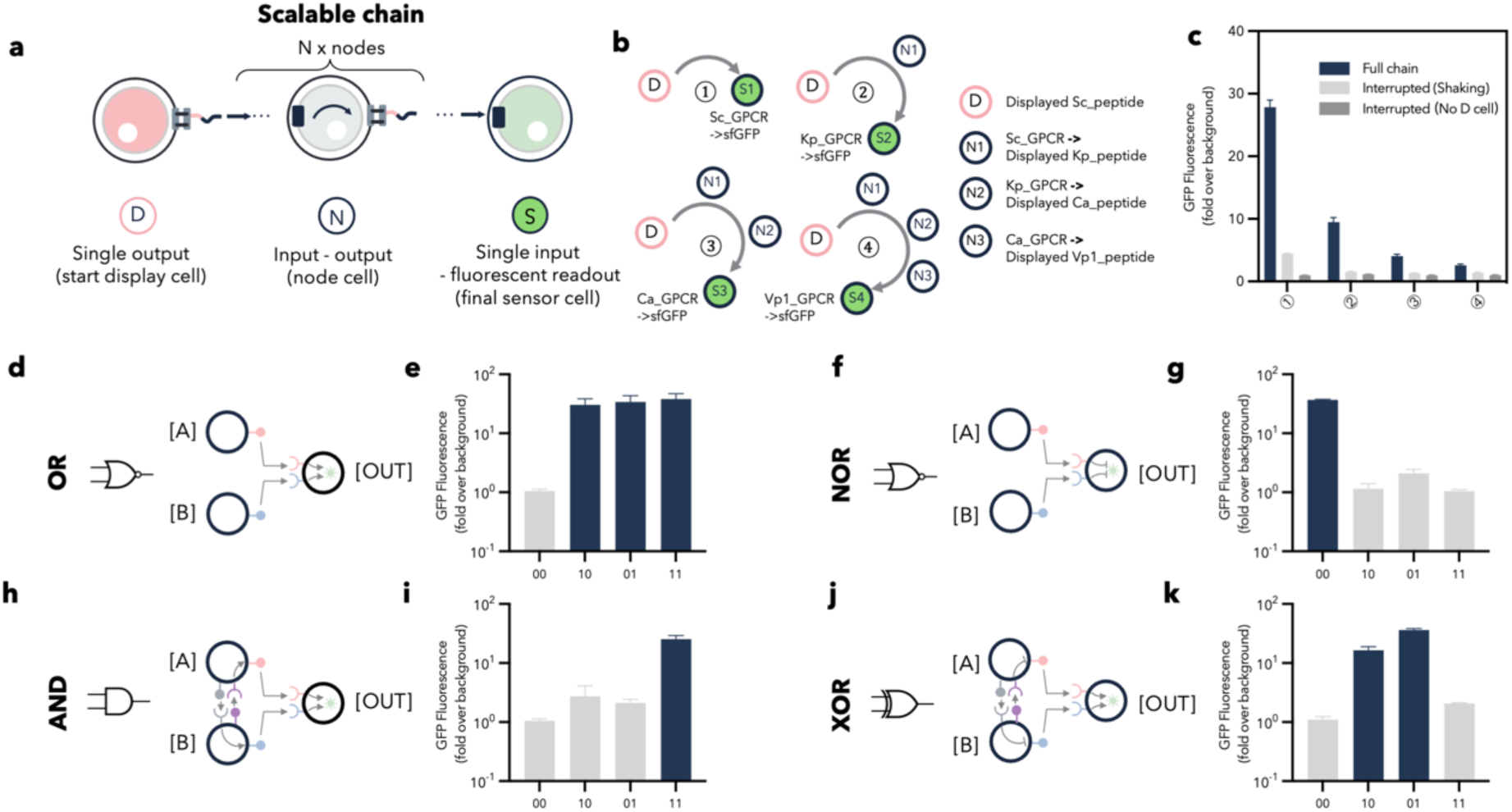
| Design of multicellular logic using MARS. **a-c** Schematic representations (**a, b**) and characterisation (**c**) of a scalable chain involving 0-3 nodes. Data for the full chain and interrupted (no D cell) configurations were recorded under static conditions. The interrupted (shaking) configuration data were recorded under shaking conditions (700 RPM); all data were captured at the 4-hour time point. **d-k** Design and characterization of OR (**d, e**), NOR (**f, g**), AND (**h, i**) and XOR (**j, k**) logic gates. ’00’ represents just sensor cells; ’10’ represents a co-culture of [A] and [OUT]; ’01’ represents a co-culture of [B] and [OUT]; ’11’ represents a co-culture of [A], [B], and [OUT]. All data show the sfGFP expression from the [OUT] cell and were recorded at the 4-hour time point except for **k**. For **k**, [A] and [B] were initially co-cultured for 6 hours under static conditions, followed by the addition of [OUT] to a 10-fold diluted [A]-[B] co-culture, with [OUT] cell sfGFP data recorded after a further 4 hours of co-culture. Experiments were conducted in triplicate and error bars show standard deviation.

Logic plays a major role in multicellular synthetic biology, and can be used for detecting cancer signals or for regulating production in co-cultures with distributed metabolic pathways^29–32^. We reasoned that by distributing different genetic logic circuits in different cells, we can leverage MARS to regulate multicellular systems. Thus, we devised MARS-based designs to create cells that perform four types of logic as a consortium. In our design for an OR logic gate the presence of either displayed Sc_peptide on the A cell or displayed Kp_peptide on the B cell suffices to activate the sensor cell (labelled as OUT) with both Sc and Kp_GPCRs (**Fig. 2d, m and Fig. S6a**). And then by incorporating a TetR-pTet-based NOT gate in this OUT cell, the OR logic design can be inverted to achieve NOR logic (**Fig. 2f, g** and **Fig. S6b**). For an AND logic gate, we established additional interactions between A and B cells that determine whether they act on the OUT cell. The A cell constitutively displays Vp1_peptide and has the Ca_GPCR, while the B cell constitutively displays Ca_peptide and has the Vp1_GPCR. In the presence of both A and B cells, contact signalling induces expression and display of Sc_peptide in A cells and Kp_peptide in B cells. These then activate the OUT cells meaning circuit output is only achieved if A and B cells are in contact (**Fig. 2h, i and Fig. S6c**). For XOR logic, expression and display of the Sc and Kp peptides in response to A and B cell contact was inverted by adding a TetR-pTet-based NOT gate into each cell. In this design OUT cells are activated when either A or B cells are present but not if both are in contact (**Fig. 2j, k and Fig. S6d**).

### SATURN, a synthetic cell-cell adhesion toolkit

One of the essential properties of multicellularity is cell-to-cell adhesion, which is especially powerful for achieving complex systems and patterning when sets of cells are designed to selectively bind each another, while excluding others^14,33–36^. While *S. cerevisiae* cells can naturally adhere to one another to take multicellular states, such in flocculation, the cell surface proteins that mediate this are not selective. Therefore, to add specific cell-to-cell adhesion as a controllable property for yeast, we developed a toolkit termed SATURN (**S**accharomyces **A**dhesion **T**oolkit for m**U**lticellular patte**RN**ing) designed for orthogonal and programable adhesion that operates independently of yeast’s own intrinsic adhesion and aggregation mechanisms.

Our design leverages yeast surface display to place high affinity binding proteins on the surface of engineered cells that specifically pair with proteins displayed on the surface of other cells (**Fig. 3a**). For this, we selected a variety of protein pairs from diverse sources known for their mutual binding properties, including nanobody-antigen pairs, dockerin-cohesin, spyTag-spyCatcher, and coiled-coil peptides. Nanobodies are small antibody fragments that have been expressed on the surface of many cell types, including yeast, and have been used in *Escherichia coli* as adhesion modules for multicellular patterning^33^. Cohesins are the main building blocks of scaffoldins that organise cellulolytic subunits with dockerin domains into multienzyme complexes^37^ and have high affinity interaction and species-specificity^38^. SpyTag-spyCatcher is a ‘molecular glue’, an irreversible spontaneous binding system between two protein domains that forms an intermolecular isopeptide bond between them^39^. Coiled-coils (CC) are well-studied protein domains that bind one another via α-helical secondary structures and have been used for cell-cell adhesion in mammalian cells^35^.

**Fig. 3.**
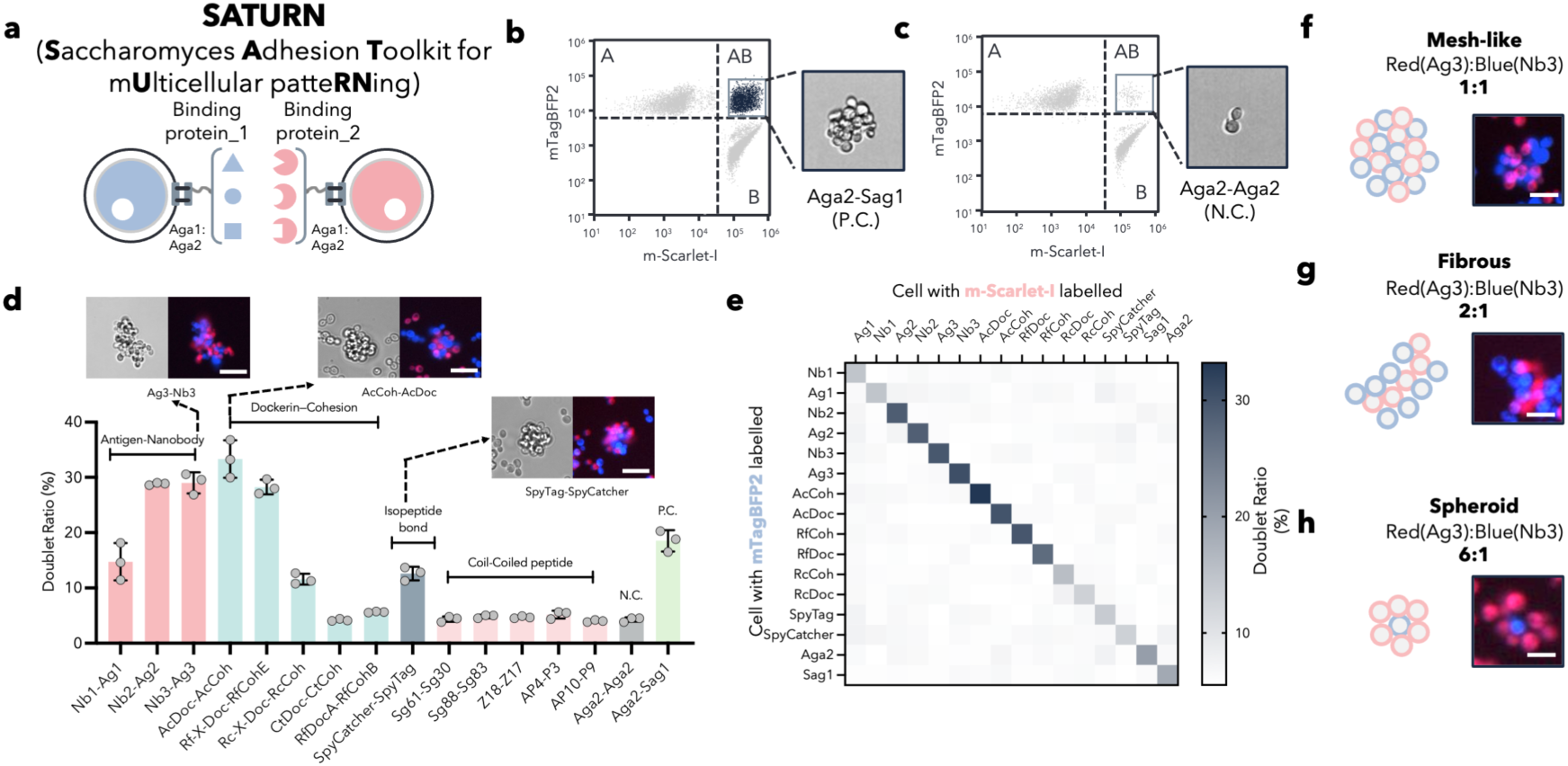
| Development of the SATURN toolkit enabling synthetic cell-cell adhesion. **a** Schematic of the synthetic adhesion design utilising Aga1-Aga2 yeast surface display technology, enabling aggregation of blue cells (B, labelled with mTagBFP2) and red cells (R, labelled with mScarlet-I) through protein-protein interactions via binding proteins. **b,c** The two-colour adhesion assay quantifies the binding strength between cells displaying Aga2-Sag1 (**b**, P.C., positive control) and Aga2-Aga2 (**c**, N.C., negative control) by measuring the doublet ratio of the whole population using flow cytometry. **d** Characterisation of the doublet ratio for B and R cells displayed with 14 different adhesive protein pairs; Scale bars in microscopy images represent 50 μm. **e** Characterisation of the orthogonality among B and R cell groups displayed with 14 adhesive protein pairs; data were recorded at the 6-hour time point under static conditions using 96 deep-well plate. **f-h** Microscopy images illustrating simple patterns achieved by adjusting the ratio of cells displaying Ag3-Nb3. Scale bars are 40 μm for **f** and 25 μm for **g, h**. All data were recorded at the 6-hour time point under static conditions in 96 deep-well plate. Experiments were conducted in triplicate, and error bars show standard deviation.

To create SATURN, we first decided to establish a new base yeast strain lacking native adhesion-related genes that may interfere with synthetic orthogonal adhesion and adhesion-driven selection. On top of the 17 gene knockouts in the strain developed for MARS, we performed a further 6 CRISPR-mediated gene deletions to remove remaining adhesins and the FLO genes, which are involved in stress-triggered flocculation, culminating in the strain yWS3516 (**Fig. S7** and **Table S1**).

Using this strain we next developed an assay utilizing flow cytometry to evaluate the efficacy of protein pairs for mediating cell adhesion (**Fig. 3b, c**). The assay uses cells genetically labelled to give red (R, mScarlet-I) and blue (B, mTagBFP2) fluorescence, with each displaying different binding pair proteins. Aggregation of these cells, triggered by the interactions of the binding proteins, results in their detection in flow cytometry as a single event giving both R+B fluorescence; an event we refer to as a ‘doublet’. The stronger the adhesion, the higher the doublet ratio detected. Through systematic comparison of the doublet ratios for various protein combinations in this yeast assay, we identified eight pairs capable of cell adhesion: Nb1-Ag1, Nb2-Ag2, Nb3-Ag3, AcDoc-AcCoh, Rf-X-Doc-RfCohE, Rc-X-Doc-RcCoh, SpyCatcher-SpyTag, and the native yeast adhesion pair Aga2-Sag1. Microscopy of sample pairs with a high doublet ratio confirmed that they indeed caused significant aggregates, typically forming clusters of more than 20 cells (**Fig. 3d**, top). Finally, orthogonality among these protein pairs was tested using the same assay, and this confirmed a remarkable level of specificity effectively ensuring that each pair functions without cross-interactions (**Fig. 3e**).

To assess whether these adhesive pairs have the potential to drive simple patterns, we tested whether multicellular morphology could be controlled by varying the ratio of the adhesive cell types. By adjusting proportions of cells in combinations, we observed changes in patterns visible by microscopy, including mesh-like structures (R:B ratio 1:1), elongated fibrous forms (R:B ratio 2:1) and spheroid structures (R:B ratio 6:1) (**Fig. 3f-h**). It is important to note that although we continuously observe such patterns by microscopy, the images selected here are the best representatives. We consider this preliminary work towards demonstrating the potential of SATURN to generate multicellular patterns in unicellular yeast.

### Combining MARS and SATURN for advanced multicellular interactions

Having established toolkit for synthetic adhesion and close-contact cell-to-cell signalling, we next set out to combine these in yeast. We first tested the effect of adhesion pair Ag3-Nb3 from SATURN on its ability to mediate MARS signalling (**Fig. S8a**) and our results showed that Vp1_MARS with Ag3-Nb3 adhesion between cells gave a higher output level than a no-adhesion control (**Fig. S8b**). Given the orthogonality in SATURN, this suggests that MARS signalling could be further controlled by engineering specific adhesion interactions between cells, giving a route for increasing complexity in multicellular behaviours.

To demonstrate the potential for these tools to allow complex patterns in yeast communities, we focused on the design of synthetic populations with three different cell types that combine MARS and SATURN. For each pattern, we first established how SATURN mediates the interactions in the population, and then add MARS so that these interactions lead to gene expression changes happening in defined cells, as they would in a natural developmental program that leads to cell differentiation.

The first pattern, ’Isolation’, uses three cell types each labelled by distinct fluorescent proteins and SATURN directing cells only adhere to those of the same type (**Fig. 4a**). Our flow cytometry assay confirmed this, showing that all events with two fluorescent colours (doublet) or three colours (triplet) were rare (**Fig. 4b**), and this was visually confirmed by microscopy (**Fig. 4b, bottom right**). When we then added Vp1_MARS signalling to these cells, so that one type displays Vp1_peptide and the other two types carry the sensor for this and produce fluorescent proteins when activated by close contact. In this cell population, with SATURN and MARS combined, we observed that the display cells were ineffective in activating the other two types of cells, presumably due to insufficient contact (**Fig. 4c, d**).

**Fig. 4.**
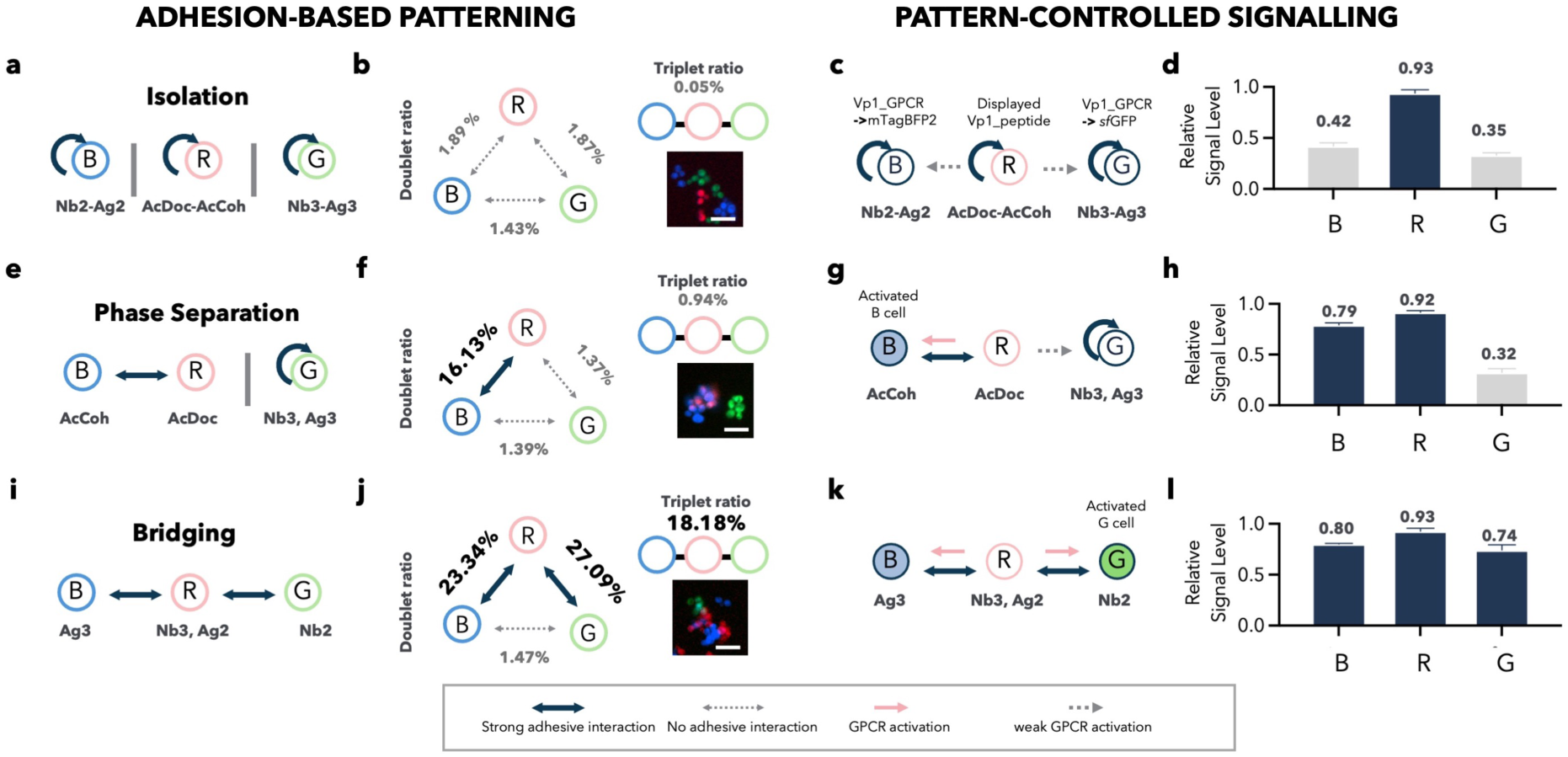
| Adhesion-mediated gene expression within a multicellular community. **a,b** Design (**a**) and characterisation (**b**) of the Isolation pattern, where the absence of adhesion leads to the isolation of three distinct cell types (R, G, B) in the adhesion assay, as quantified by the % doublet values annotating the interaction arrows and the % triplet ratio shown above the microscopy example (scale bar = 50 μm). **c,d** Upon incorporation of MARS, the Isolation pattern prevents effective activation of sensors in G and B cells from ligand display by the R cell due to their separation. **e,f** Design (**e**) and characterisation (**f**) of the Phase Separation pattern. Self-adhesive homophilic (Ag3/Nb3/sfGFP) and heterophilic (AcCoh/mTagBFP2 + AcDoc/mScarlet-I) pairs result in a distinct phase separation phenotype. **g,h** When combined with MARS, the Phase Separation pattern means that the displaying R cell effectively activates the B sensor cell but not the non-interacting G sensor S2 cell. **I,j** Design (**i**) and characterisation (**j**) of the Bridging pattern. The presence of a doubly adhesive strain (Nb3/Ag3/mScarlet-I) enables a pattern with bridging between non-interacting cells (Ag3/mTagBFP2 and Nb2/sfGFP). **k,l** The Bridging pattern enables the R display cell to activate both the B and G sensor cells. Experiments were done in triplicate and errors represent the SD. The method for calculating relative signal levels is given in the methods section. Data for **b, f** and **g** were taken at the 6-hour time point under static growth conditions in 96 deep well plates. **d, h** and **I** were taken at the 4-hour time point under 250 RPM shaking conditions in 96 deep well plates. Cocultures of **c, g** and **k** at static condition or high-speed shaking conditions (700 RPM) eliminated any difference shown in **d, h, l** (data not shown).

In the next pattern, ’Phase separation’, SATURN was used to allow cells labelled with mTagBFP2 (B) and mScarlet-I (R) to cluster through AcCoh-AcDoc interactions, while cells labelled with sfGFP (G) were excluded, instead adhering homophilically via simultaneous display of Ag3 and Nb3 (**Fig. 4e**). In this population we saw a big increase in the occurrence of B+R doublet events (>16%), whereas all possible doublet or triplet events involving G cells were rare (**Fig. 4f**) and microscopy analysis confirmed this pattern (**Fig. 4f, bottom right**). Adding Vp1_MARS signalling to these cells now gave a system where display cells could selectively activate only one of the two types of sensor cells when mixed in the population, with the contact between these cells mediated by the adhesive protein pair (**Fig. 4g, h**).

The last pattern, ’Bridging’ uses SATURN so that R cells display both Nb3 and Ag2 and adhere to both B cells expressing Ag3 and G cells expressing Nb2. This promotes a unified phase among all three cell types with the adhesion assay showing a big increase in events that are R cell-involving doublet (>50%) or triplets (>18%) (**Fig. 4i**). Doublets involving only B and C cells stay rare in this configuration (**Fig. 4j**) and microscopy confirms that R cells aggregate with B and C cells effectively acting as a bridge to connect B and G cells (**Fig. 4j, bottom right**). Adding Vp1_MARS signalling to these cells, so that the bridging cells display the peptide and the other cells are sensors, leads to R cells being able activate gene expression in both the B and G cells in the patterns (**Fig. 4k, l**). These three patterns showcased here demonstrate the capabilities for engineering sophisticated interactions within a multicellular *S. cerevisiae* community using MARS and SATURN together.

### JUPITER, a yeast system for monitoring protein-protein interactions

Monitoring protein-protein interactions (PPIs) such as nanobody-antigen interactions is valuable for drug discovery, tool development and basic research. Yeast and bacterial two-hybrid systems have long been effective tools for mapping PPIs^40^, but these focus on assessing interactions of intracellular proteins. Surface-display technologies, on the other hand, facilitate exploration of extracellular PPIs in customizable environments and have been extensively employed to isolate novel antigen-specific antibodies with therapeutic applications^41^. While mammalian surface display of antibodies preserves critical native features of these molecules, such as glycosylation patterns, only limited sizes of libraries is achievable with such cells. Thus, yeast surface display (YSD), which can exploit the ease of making and screening very large libraries with yeast, has become a prevalent technique for screening antibody libraries^41^ and has led to the development of genetic systems in yeast that characterize PPIs by tricks such as reprogramming yeast mating and coupling interaction events with next-generation sequencing^17,42^. Despite its potential, most YSD-based screening methods involve interactions only at the cell surface and either cannot link to triggering genetic responses in the cells or the genetic responses require the cells to commit to a mating cycle. A system where PPIs directly trigger gene expression in the cells during their growth and proliferation would be especially valuable for long-running experiments, such as those involving *in vivo* continuous evolution and selection. To this end, we combined MARS and SATURN to develop JUPITER (Juxtacrine sensor for Protein-protein InTERaction), a system that links the binding strength between two extracellular proteins of interest (POIs) to intracellular gene expression responses. The design of JUPITER is modular, allowing users to display one POI, such as a nanobody, via yeast surface display system on the sensor cell of MARS system. Then in the display cell, the corresponding MARS peptide ligand and another POI, such as an antigen, are co-displayed (**Fig. 5a, left**).

**Fig. 5.**
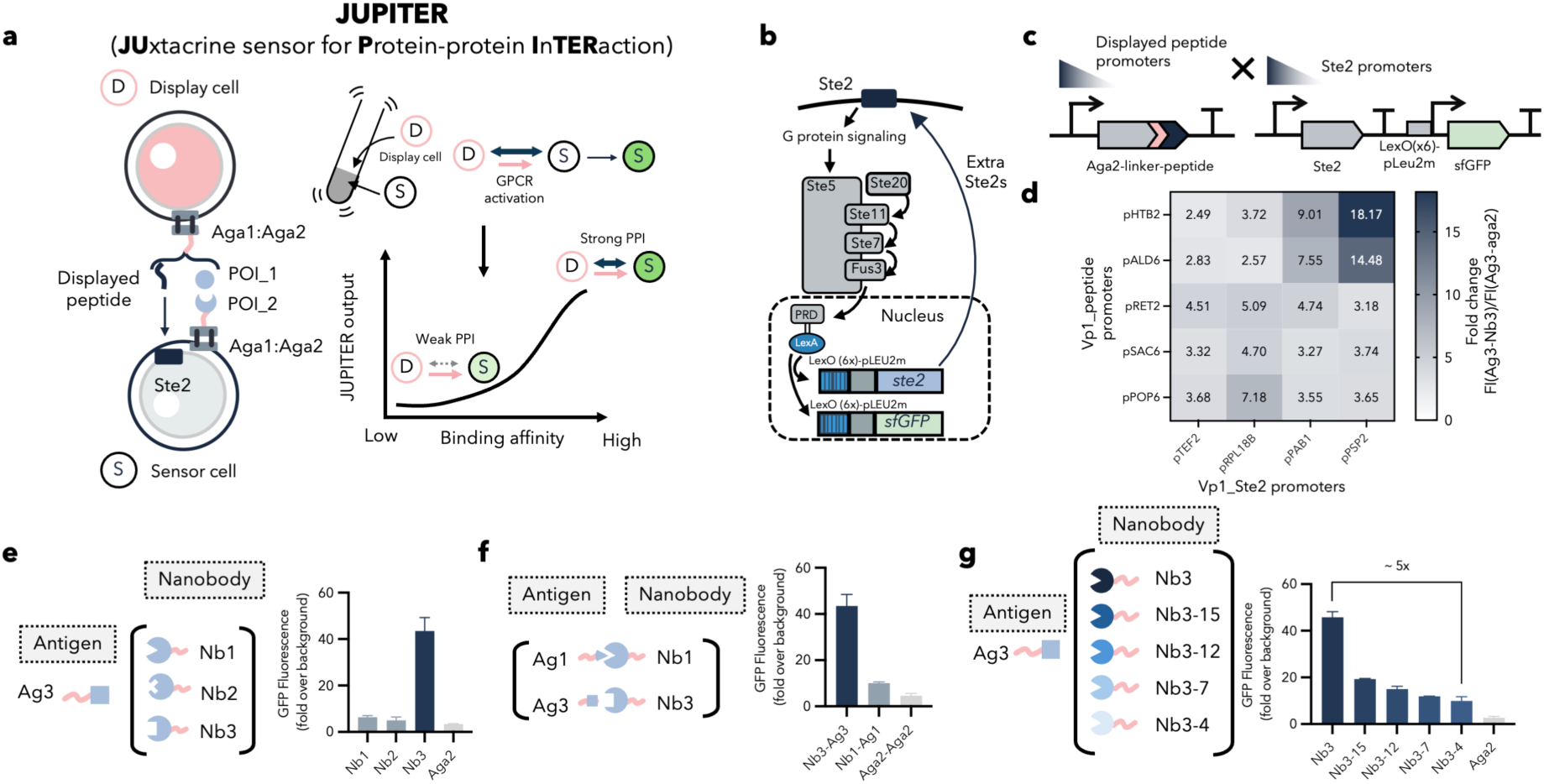
| Development of JUPITER for testing protein-protein interactions. **a** Schematic of the JUPITER design, functioning as a genetic sensor for monitoring protein-protein interactions between display (D) and sensor (S) cells. **b** An additional Ste2 GPCR expression cassette was integrated into the yeast with a signalling-controlled promoter that drives extra production of Vp1_GPCRs on the cell membrane in response to ligand sensing. **c,d** A variety of promoter combinations were tested to fine-tune the expression levels of the displayed peptide and GPCR sensor for JUPITER. Optimal promoter combinations, pHTB2-Vp1_peptide and pPSP2-Vp1_Ste2 were used for all later JUPITER experiments. **e** JUPITER specifically activates sfGFP expression in sensor cells displaying the nanobodies (Nb) with the highest affinity to the antigen (Ag3) displayed on the display cell. **f** JUPITER is then utilised to compare the relative binding capabilities of various nanobodies, with protein-protein affinities correlating to sensor cell activation levels. **g** JUPITER is capable of differentiating between different variants of the same nanobody. All data were captured at the 7.5-hour time point under shaking conditions (700 RPM) using 96 deep well plate. Experimental measurements of sfGFP levels per cell were determined via flow cytometry and are presented as the mean ± standard deviation from three replicates.

We anticipated that in shaking conditions, if no cell adhesion is mediated by POI-POI interactions, then the opportunities for the MARS GPCR to contact its displayed peptide would be reduced, preventing effective activation of the sensor cell type. Conversely, when interaction strength between target proteins is high, contact opportunities between sensor and display cells are increased, enabling sensor cells to be activated even in a shaking condition (**Fig. 5a, right**). As all components of JUPITER are based on the modular MARS and SATURN toolkits, all aspects of the set-up, from the POI display strength to the downstream signalling response, can be customised and fine-tuned for each application.

To establish JUPITER we tested a version where display and sensor cells detect and respond to Ag3-Nb3 interactions. However, we first needed to optimise MARS as we observed that it operated with a high background when SATURN components are added (**Fig. S8b)**. To reduce the background, we integrated a feedback mechanism in the MARS sensor cells by incorporating an additional Ste2 GPCR gene cassette in the genome whose expression is itself activated by MARS signalling. Activation of MARS leads to increased production of GPCR in the sensors, enhancing contact opportunities with the peptide of the display cells and this intensifies the downstream signal (**Fig. 5b**). Then by optimising expression levels of Vp1_Ste2 GPCR and Vp1_peptide, we found promoter combinations that optimize the dynamic range of JUPITER from less than 2-fold to more than 18-fold when Ag3-Nb3 interactions occur (**Fig. 5c, d**).

The optimised JUPITER system was then shown to demonstrate excellent orthogonality for distinct protein-protein interactions from our SATURN toolkit; with only the correct antigen-nanobody pairs capable of triggering strong gene expression output in sensor cells (**Fig. 5e**). JUPITER’s ability to distinguish between specific PPIs makes it a valuable tool for studying protein interactions in complex biological systems. Importantly, in our experiments JUPITER accurately reflected the known relative strengths of the different PPIs we tested, with the sensor cell expression output concordant with the interaction strengths of Nb3-Ag3, Nb1-Ag1, and aga2-aga2 pairs (**Fig. 5f** vs. **Fig 3d**). Further investigation into the limits of JUPITER revealed its ability to distinguish between variants of the same nanobody, with fluorescent protein expression levels in sensor cells varying over 5-fold from when the weakest nanobody for the Ag3 antigen was used (Nb3-4), compared to when the strongest antibody was used (Nb3), as is consistent with reported data for these proteins^33^ (**Fig. 5g**). These encouraging results highlight JUPITER’s potential for studying PPIs in yeast cultures.

### Applying JUPITER for targeted nanobody screening and selection

We anticipate that JUPITER can serve as a screening system for assessing and selecting protein pairs with enhanced binding capabilities. A major advantage of JUPITER over other PPI assay systems is that it is a genetic sensor and so a positive interaction can be readily linked to custom downstream outputs. As an initial validation of JUPITER, we examined if sensor cell output could be used to induce observable phenotypic differences in cells, such as altered growth rates. To this end we engineered a fusion protein, His3-2A-sfGFP, to be the output gene, with this giving simultaneous production of the auxotrophic selection enzyme His3 and sfGFP in activated yeast cells through the self-cleaving action of the 2A peptide upon expression (**Fig. 6a**). We evaluated a JUPITER set-up using this output gene by assessing growth in minimal media with varying concentrations of histidine. Our findings indicated that reducing the histidine concentration to 50% of the normal levels allows JUPITER to effectively balance fluorescence intensity and cell growth (**Fig. 6b, c**). This 50% histidine concentration was established as the standard condition for subsequent screening experiments where activated output should lead to improved cell growth.

**Fig. 6.**
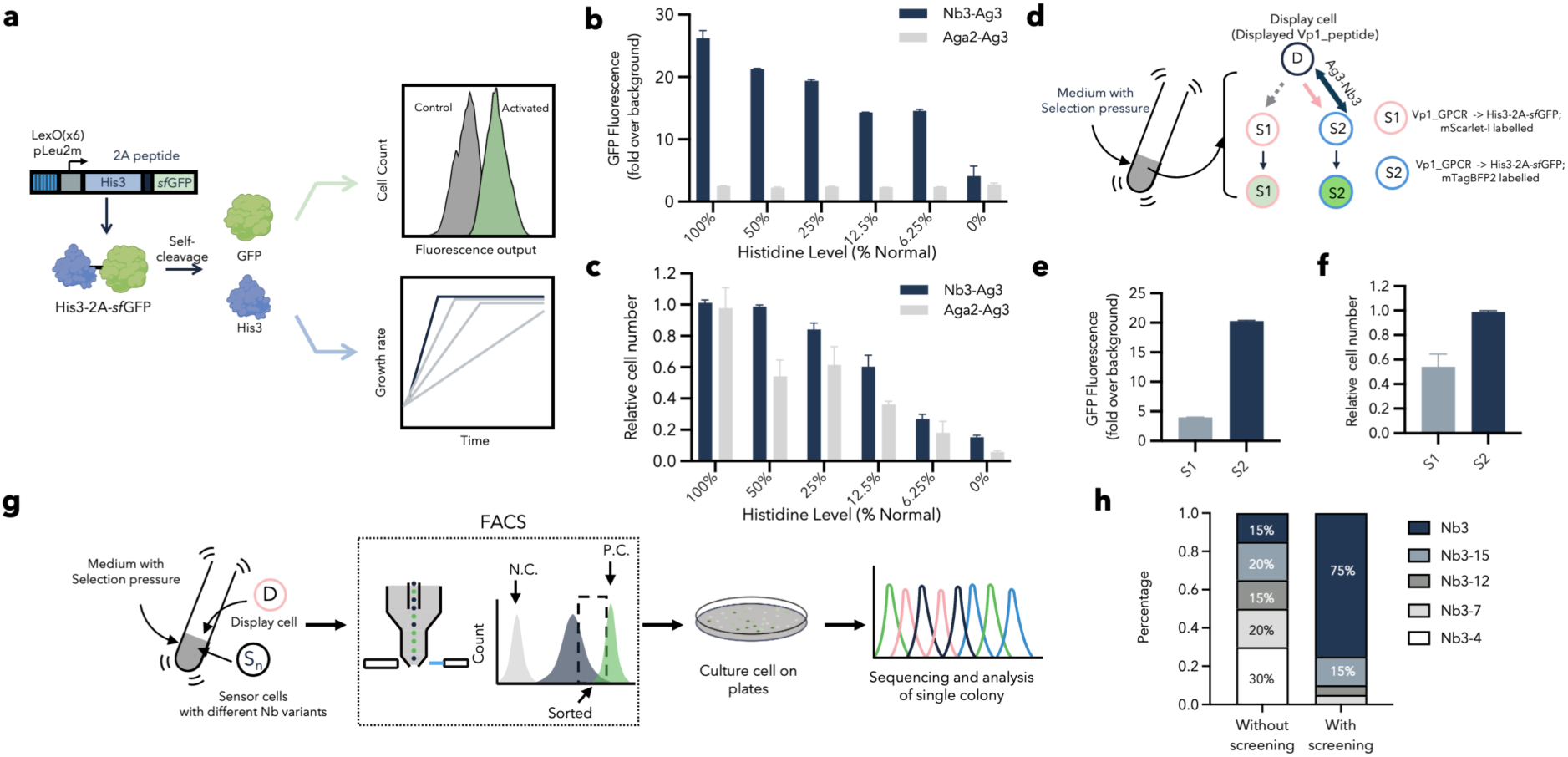
| Applying JUPITER for targeted nanobody screening and selection. **a** His3-2A-GFP fusion protein enables the simultaneous production of His3 and sfGFP via the self-cleaving action of the 2A peptide upon expression. This leads to improved growth in reduced histidine media and increased fluorescent output when expressed in activated sensor cells. **b,c** Reducing histidine media concentration to 50% of standard level allows JUPITER to effectively differentiate sensor cells by their fluorescence intensity (**b**) and cell growth (**c**). **d-f** Interaction between the S1 cell and the D cell results in higher cell numbers and fluorescence output compared to the non-interacting S2 cell during JUPITER screening. **g** Schematic representation of JUPITER applied using FACS for screening a library of nanobody variants. Sensor cells whose sfGFP fluorescence falls between those of negative control (N.C.) and positive control (P.C.) are sorted, cultured and their DNA sequenced. **h** Sanger sequencing results from before or after screening a library of five Nb3 variant displaying sensor cells show the relative levels of the different cells in population, based on 20 colonies being analysed for each group. All fluorescence data was recorded at 7.5-hour time point in 96 deep-well plate in shaking conditions (700 RPM). Experimental measurements are sfGFP levels per cell determined by flow cytometry and shown as the mean ± SD from triplicate isolates.

Next, we evaluated the ability of JUPITER to differentiate 2 different sensor cells in a mixed population, in this case ones displaying the high-affinity Nb3 nanobody versus those only displaying aga2 and no nanobody (**Fig. 6d**). In this setup, S1 sensor cells, tagged with mTagBFP2 and displaying the Nb3 protein, were expected to interact with a display (D) cell presenting Ag3 antigen. Conversely, S2 cells, labelled with mScarlet-I and serving as the control group should not bind well to D cells. After co-culturing these three types of cells in selective media, flow cytometry was used to measure GPCR-induced fluorescence output (sfGFP) in S1 and S2 cells and compare the relative number of S1 and S2 cells in the population. After 7.5 hours of co-culture, results showed that S1 cells had stronger GFP expression and were found in higher numbers compared to S2 cells, validating the approach (**Fig. 6e, f**). We then took this further, combining the assay with flow cytometry sorting (FACS) to screen and select high affinity binders from a pool of yeasts displaying various Nb variants with different affinities to Ag3 (**Fig. 6g**). This screening demonstrated that Nb3, the nanobody with the strongest reported binding affinity to Ag3, was found on 75% of the yeast after JUPITER screening and cell sorting (**Fig. 6h**). These proof-of-concept experiments demonstrated the feasibility of employing JUPITER in genetic screening and selection systems, offering a promising avenue for future applications including enhancing desired protein-protein interaction through *in vivo* directed evolution.

## Discussion

In this work, we introduced MARS as a toolkit to achieve cell-cell contact-dependent signalling in *S. cerevisiae*. This was achieved by modifying the traditional secretion pathways to display mating peptides from fungi directly on the cell surface, altering the communication mode between cells. Given the variety of natural signalling processes that utilise secreted peptides, this approach can potentially be adapted to other organisms including bacteria^43,44^, viruses^45,46^, mammalian cells^47,48^, and so expand the range of applications for cell-cell contact-dependent signalling across different biological kingdoms. However, the complexity of peptide-mediated interactions, with peptides activating receptors in many different ways, underscores the necessity of refining display methods. It is essential to ensure that displayed peptides both expose their receptor-binding domains effectively from the cell surface, but also retain the correct structure to effectively activate receptors.

The contact-dependent activation enabled by MARS mimics juxtacrine signalling, however we stop short of calling it juxtacrine here, as direct evidence of displayed peptide binding to its receptor on another cell is not provided. The need for charged amino acids in the linker of the displayed peptide ligand also hints that other mechanisms may be involved, yet despite this uncertainty, MARS can effectively recreate juxtacrine signalling in all our assays and applications.

For multicellularity, cell-to-cell adhesion is of equivalent importance to signalling, and our modular synthetic adhesion toolkit SATURN provides for the first time a system for programming *S. cerevisiae* cells to selectively bind in pairs and sets using eight genetically encoded, orthogonal protein pairs. We envision wide use of SATURN for experiments and applications where yeast multicellularity is required. When combined with SATURN, we show that MARS can direct yeast cells to alter gene expression in response to multicellular environments, mimicking how cells in multicellular systems differentiate in response to their surrounding cells. To our knowledge, MARS and SATURN together represent the first demonstration of synthetic juxtacrine-like signalling and adhesion systems in yeast and expand the possibilities for engineering in this common synthetic biology chassis. The two toolkits have the potential to integrate seamlessly with existing and emerging synthetic biology tools for yeast, opening new avenues for exploring *S. cerevisiae*’s capabilities for biosensing, biomanufacturing, and for making therapeutics^49^.

Finally, we developed JUPITER, a proof-of-principle biotechnology application made possible by combining MARS and SATURN. JUPITER is novel genetic sensor system for extracellular protein-protein interactions and succeeded in screening and selecting for high-affinity nanobodies from a test pool of engineered cells. JUPITER offers several advantages over traditional yeast-display methods used for screening nanobody libraries. The system is highly modular, so that by simply replacing the target protein displayed on the sensor cell, it facilitates immediate initiation of characterization and screening of new protein-protein interactions of interest. Notably, JUPITER eliminates the need for purified antigen molecules, as these are displayed directly on the surface of the display cell, reducing complexity and cost. This attribute is particularly advantageous for initial high throughput testing experiments and may aid in the development of biomedical applications. JUPITER also allows for the customization and adjustment of the downstream output without requiring cellular or environmental changes. This makes it highly suitable for long-term use, such as with *in vivo* continuous evolution processes like Orthorep^50^. While JUPITER has been validated in smaller, focused libraries, its effectiveness for larger, more diverse protein libraries remains to be fully explored. So far, we have only demonstrated JUPITER’s ability to show the strength of interactions between protein mutants of similar size or structure. Further work is needed to determine whether JUPITER can effectively distinguish the interaction relationships between proteins that have significant size or structural differences.

In establishing these exciting multicellular engineering tools for yeast, we also developed a useful new base yeast strain with extensive genome modifications that remove 23 of yeast’s genes directly involved in its native ability to aggregate under stress and do other multicellular phenotypes (yWS3516, **Fig. S7** and **Table S1**). We envision this strain of having value for many other applications and studies. In past work exploiting and tuning GPCR signalling in yeast, we made an analogous ‘stripped-back’ yeast strain with many components of the mating signalling pathway removed by gene knockout, which has since been used in several other studies (yWS677^51–53^). It has not escaped our notice that in the work presented here we had to reintroduce feedback control into the signalling in our base strains in order to achieve the most effective multicellular behaviours.

## Materials and Methods

### Strain and media

Yeast strains discussed in this work all originate from the list provided in Table S1. For the purposes of DNA cloning and the propagation of plasmids, NEB Turbo competent *Escherichia coli* (*E. coli)*, obtained from New England Biolabs (NEB), was employed throughout. *E. coli* cultures were grown in Luria-Bertani (LB) medium. When needed, antibiotics such as ampicillin (100 μg mL-1), chloramphenicol (25 μg mL-1), kanamycin (50 μg mL-1), and spectinomycin (100 μg mL-1) were added. Yeast cells were generally cultured in a defined synthetic complete media (SC) with needed amino acid supplements.

### CRISPR/Cas9 genome engineering

A mixture for yeast transformation was prepared using 250 ng of the CRISPR/Cas plasmid and 500 ng of each PCR-amplified DNA fragment. This was combined with 10 μL of boiled salmon sperm DNA and then brought to a total volume of 64 μL with nuclease-free water. Yeast cells were transformed using the lithium acetate protocol^5^. For details on the specific gRNAs utilised in CRISPR/Cas9 genome engineering, refer to Table S6.

### Flow cytometry

Fluorescence levels in cells were quantified using a Thermo Scientific Attune NxT Flow Cytometer. The settings employed were as follows: forward scatter (FSC) at 300 V, side scatter (SSC) at 350 V, blue laser (BL1) at 500 V, and yellow laser (YL2) at 450 V. For each experiment, unless otherwise specified, 10,000 events representing the yeast cell population were selected based on forward and side scatter criteria and recorded. These events were analysed using FlowJo software v10.4, with results presented as the geometric mean of the peak heights from the relevant fluorescence channels. The reported values represent the average from three separate biological replicates. Data visualisation was performed using GraphPad Prism v10.2.3.

### GPCR biosensing assay

Display cells and sensor cells were inoculated into 600 µL of synthetic complete media and cultivated in 2.2 mL deep-well plates (96 wells) at 30°C using an Infors HT Multitron shaker set at 700 rpm overnight. The following day, cultures were normalised to an optical density (OD600) of 1, as measured by a Synergy HT Microplate Reader from BioTek. These cultures were subsequently diluted at a ratio of 1:100 into fresh media. Unless specified otherwise, the diluted cultures were incubated for 4 hours in 14 mL culture tubes, either in a static state at 30°C or under shaking conditions at 700 RPM and 30°C. For flow cytometry analysis, 200 µL of culture from each well was transferred directly into a 96-well clear, flat-bottom microplate.

### Screening the amino acid composition

To investigate whether altering two amino acids preceding the peptide could enhance activation of the displayed peptide-GPCR system, degenerate primers featuring a varied NNNNNN region (N = A, T, C, G) were ordered from IDT to mutate these residues. The resultant library was transformed into Escherichia coli. Colonies were harvested from the entire plate by overlaying 1 mL of LB medium with the appropriate antibiotics and gently mixing with a plastic spreader. Plasmid mini-preparations were then performed after a 6-hour incubation at 37°C. Following standard yeast transformation procedures, the library was integrated into the desired genomic site within the yeast. 200 colonies were selected for co-cultivation with the sensor strain. The fluorescence signal levels in the sensor strain were quantified using flow cytometry. Colonies that exhibited a positive fluorescence response were subsequently sequenced and further characterised for detailed analysis.

### Adhesion assay

Each strain was inoculated into 600 µL of synthetic complete media and cultured overnight at 30°C in 2.2 mL deep-well plates (96 wells) on an Infors HT Multitron shaker set to 700 rpm. The following day, cultures were adjusted to an OD600 of 1 using a Synergy HT Microplate Reader from BioTek, and then diluted 1:100 into fresh media. These cultures were then incubated for six hours at a constant temperature of 30°C without agitation. For measuring adhesion strength, an Attune NxT Flow Cytometer (Thermo Scientific) was used without any mixing steps, and the flow rate was set at 100 µL/min. Fluorescence data for 20,000 events were captured and analysed using FlowJo software. Cell types were differentiated using quadrant gating based on their fluorescence profiles. The proportion of sfGFP+/mScarlet-I+ doublet cells was calculated to assess the relative adhesion strength of the protein pairs. For adhesion imaging, mixed adhesion cells were cultured together for two hours before imaging using an Attune CytPix Flow Cytometer. Samples were acquired at a flow rate of 100 µL/min, and images of cells displaying both sfGFP and mScarlet-I signals were captured for further analysis.

### Relative Signal Level Calculations

For the adhesion-mediated gene expression assays (**Fig. 4**) co-cultures of a sensor cell (expressing Vp1_GPCR linked to sfGFP/mTagBFP2 and displaying Nb3) and a display cell (expressing Vp1_peptide and displaying Ag3) were used in experiments as a positive control expected to produce the maximum output from the sensor cell. Output signals from the cells in this control experiment were then used to normalise the output signal from cells in all the experimental tests.

### Fluorescence microscopy

Prepare a 1.5% (w/v) low-melt agarose solution by adding 0.15 g of agarose to 10 ml of liquid bacterial medium in a 50-ml polypropylene conical tube. Dissolve the agarose by alternately microwaving and vortexing the mixture. Allow the solution to cool for several minutes. Then, carefully transfer 1 ml of the cooled agarose solution onto a 22 mm² glass coverslip, ensuring it is placed on a level surface. Immediately place another glass coverslip on top to form an agarose gel sandwich, taking care to eliminate any air bubbles. Allow the setup to solidify at room temperature (approximately 25°C) for 30 minutes. Once solidified, introduce yeast cells onto the agarose pad by dispensing 2 µL of the yeast cell mixture. Allow the agarose pad to dry for an additional 15 minutes before gently flipping the agarose pads onto a cover glass-bottom dish, ensuring the yeast cells are sandwiched between the agarose pads and the cover glass. Seal the dish with Parafilm to prevent evaporation. Observations were made using a Nikon ECLIPSE Ti microscope. The microscope was set to a 20x magnification with configurations for bright field (BF) and fluorescent channels GFP/BFP/RFP. Image processing and analysis were conducted using Fiji software.

### Fluorescence-Activated Cell Sorting (FACS)

Sorting of yeast cells was performed using the BD FACSAria III Cell Sorter from BD Biosciences. Yeast cells were first resuspended in PBS and transferred into a 5 mL FACS tube. The cell suspension was then adjusted to the required concentration with PBS prior to sorting. Cells exhibiting higher GFP fluorescence were selectively collected into a 15 mL Falcon centrifuge tube. A portion of the sorted cells was immediately cultured in SC Leu-medium and incubated overnight at 30°C and 250 rpm until saturation was reached. The remaining cells were centrifuged at 4000 rpm for 20 minutes, resuspended in 0.25 mL of PBS, and stored in 25% glycerol at -80°C for future use.

## Supplementary Materials

**Supplementary Fig. 1.**
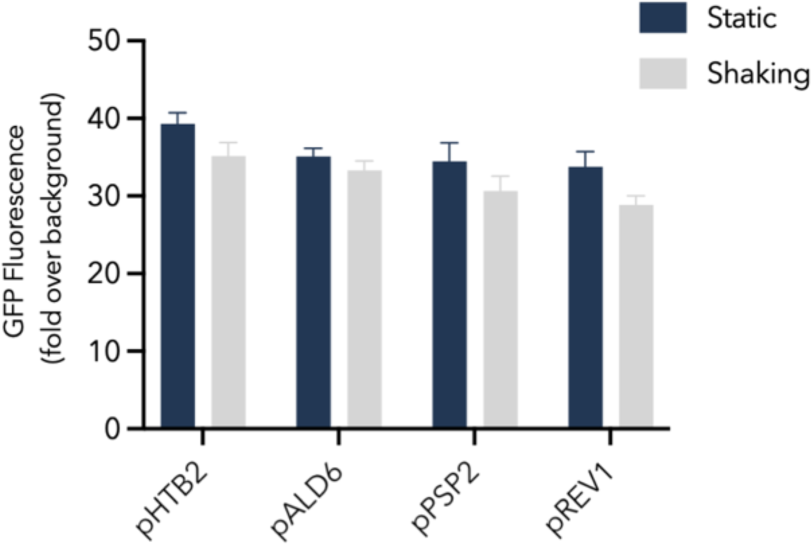
| The activation level of the Sc_GPCR sensor cell when co-culturing with a secretor cell. The secretion level of Sc_peptide was tuned using yeast promoters from high to low strength. Only a slight difference between shaking and static conditions was observed for secretor and sensor cell co-cultures. Data were recorded at the 4-hour time point. Experiments were conducted in triplicate, and error bars represent the standard deviation.

**Supplementary Fig. 2.**
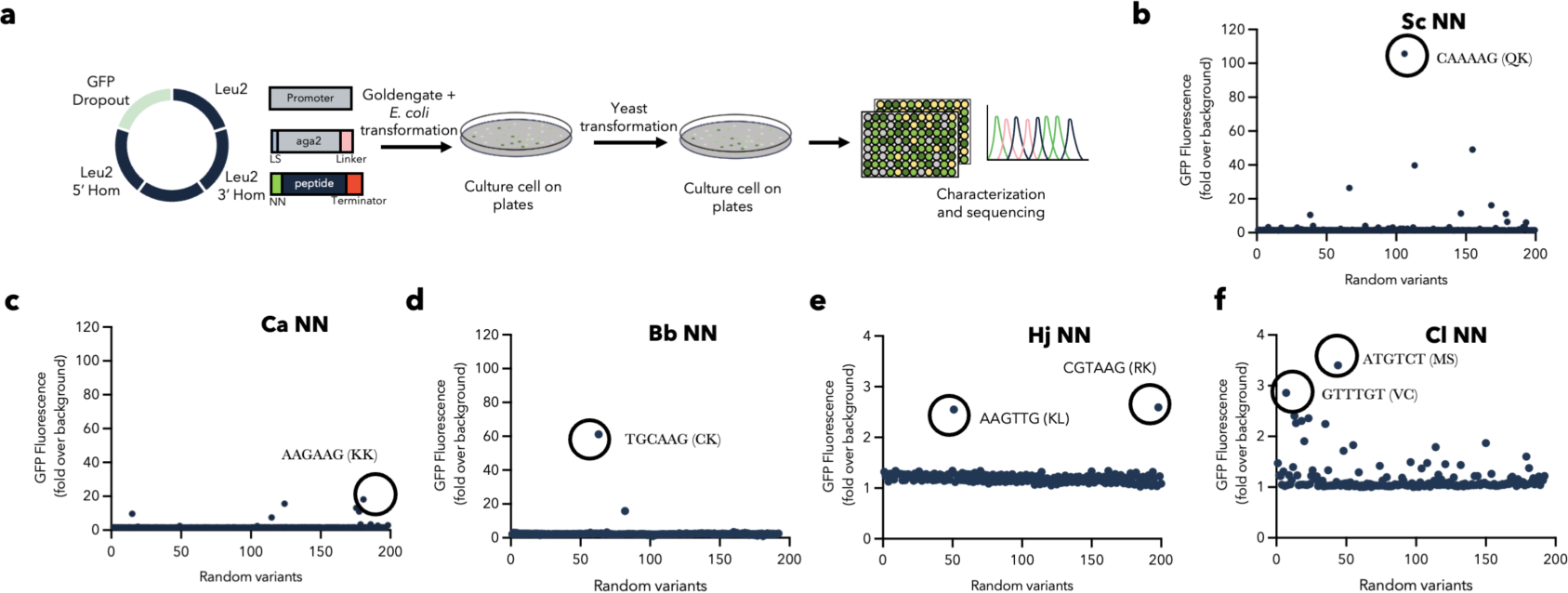
| Summary of NN strategy to develop functioning MARS ligands. a Schematic of NN strategy used to identify the optimal NN amino acids combinations for the peptide linker to be capable of activating GPCRs. b The results of GPCR activation for the colonies from Sc, Ca, Bb, Hj, and Cl, following the selection of the NN strategy. Typically, 200 colonies were picked and cocultured with the corresponding sensor cell, with the green fluorescence of the sensor determined by flow cytometry to identify promising candidates (circled, with DNA and amino acid sequence shown).

**Supplementary Fig. 3.**
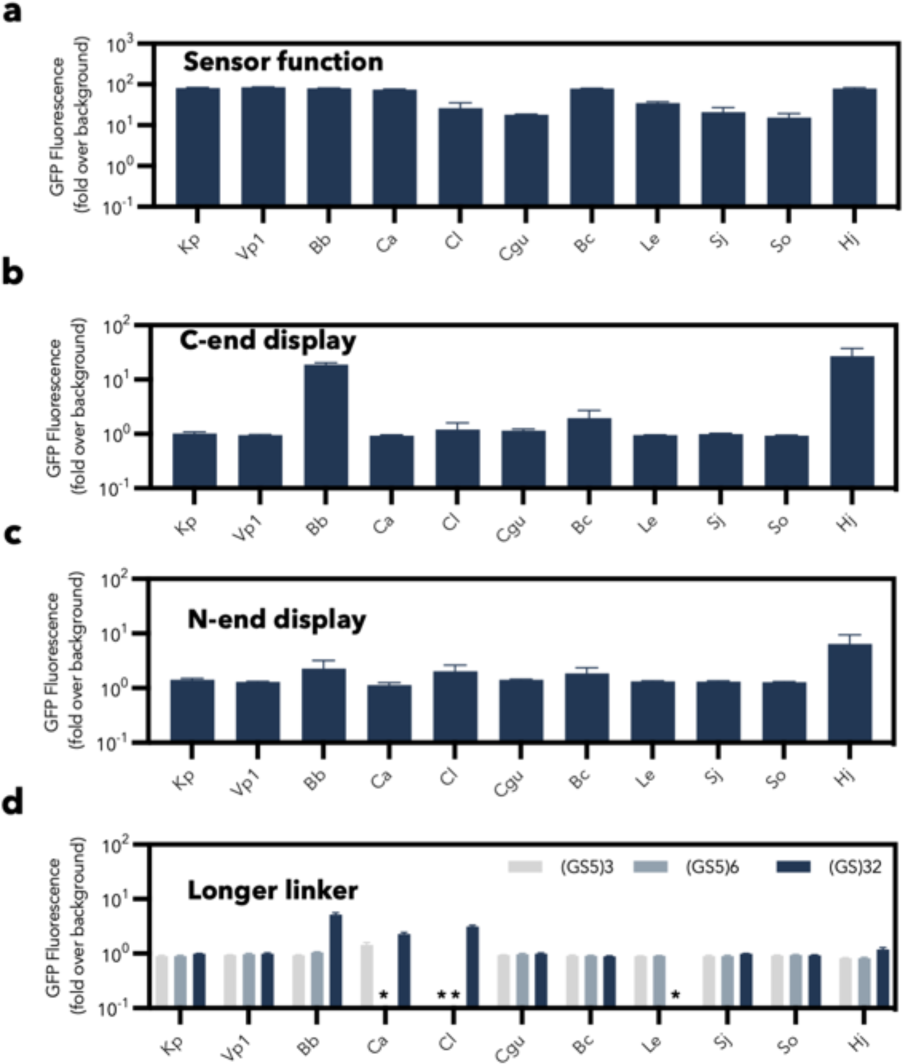
| Summary of C-end, N-end and Longer linker strategies to develop a library of synthetic functional MARSs. a All GPCRs selected for MARS worked as sensors in strain yWS2064 in terms of being detected when secreted by another cell. b-d The results of C-end, N-end and Longer linker strategies to develop more functional MARSs. Sensor cell GFP expression data were recorded at the 4-hour time point and showed no functioning activation in most cases, except for 2 promising results for C-end display. Experiments were conducted in triplicate, and error bars represent the standard deviation.

**Supplementary Fig. 4.**
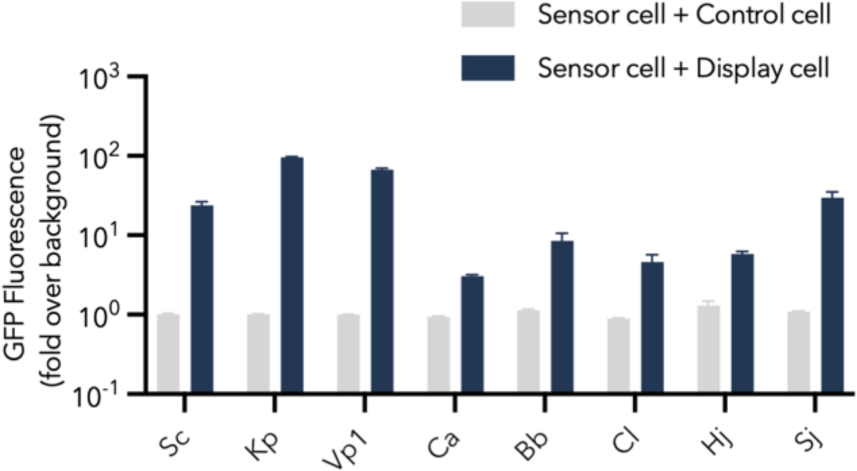
| The activation level of all MARSs developed in this work. A total of eight MARSs with cell-to-cell contact-based signalling were achieved. These showed a wide range of sensor cell activation profiles. Data were recorded at the 4-hour time point. Experiments were conducted in triplicate, and error bars represent the standard deviation.

**Supplementary Fig. 5.**
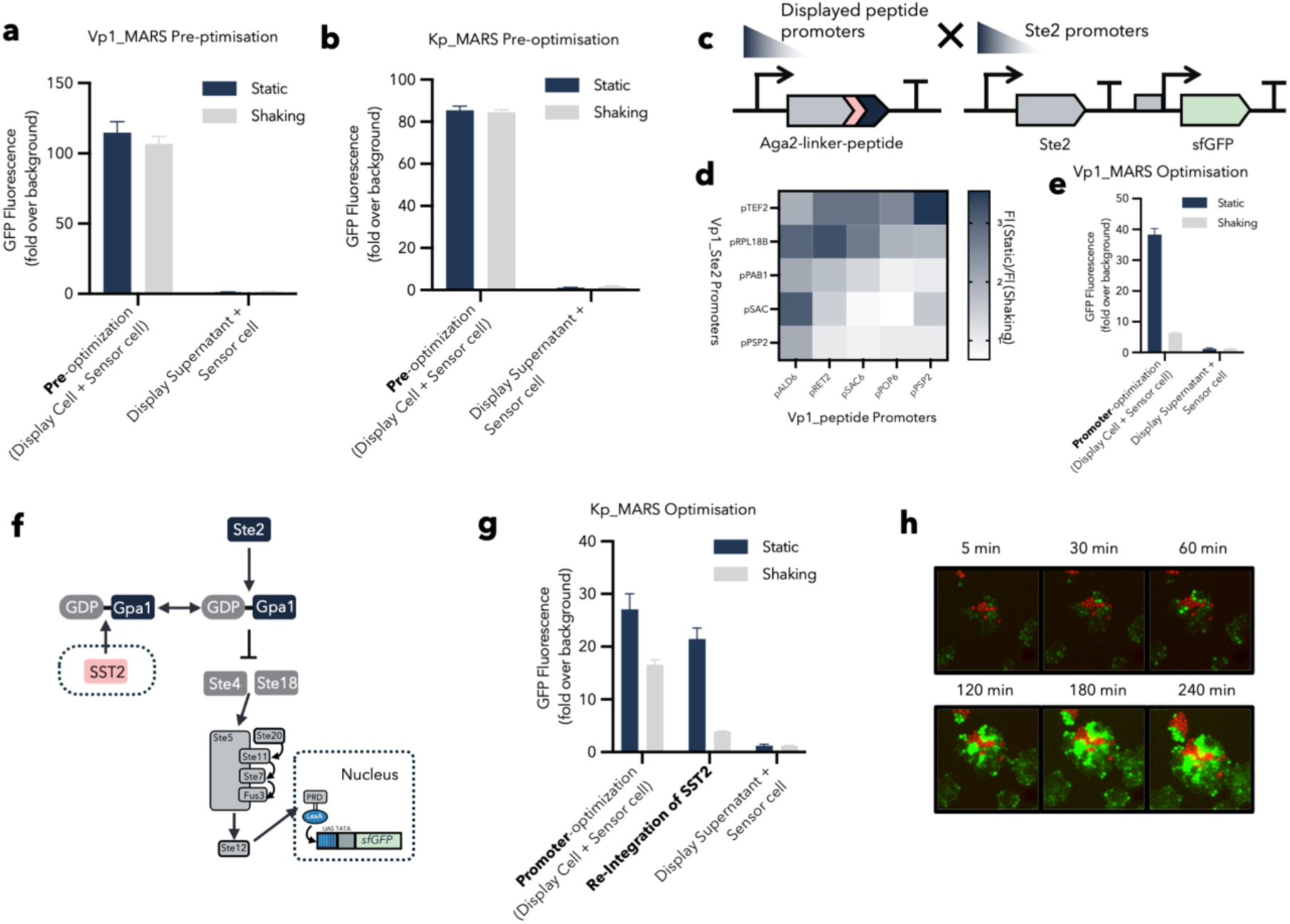
| Optimisation of Vp1 and Kp MARS systems. a,b Characterization of the Vp1_MARS (a) and Kp_MARS (b) systems before rounds of optimization. c A variety of promoter combinations were employed to tune the expression level of the displayed peptide and Ste2 GPCRs in the cells. d Heatmap of the dynamic range of Vp1_MARS between shaking and static conditions, when the system is tuned by varying the levels of expression of the Vp1_peptide and Vp1_GPCR. e Characterization of the optimized Vp1_MARS after selecting the correct promoter combinations for component expression. f Schematic showing that SST2 functions as a negative regulator within the GPCR pathway. g Characterisation of the optimized Kp MARS achieved after optimizing promoter selection for components and reintegrating SST2. h Microscope images of Kp MARS-based activation over time in a growing culture. All MARS activation data were recorded at the 4-hour time point. Experiments were conducted in triplicate, and error bars represent the standard deviation.

**Supplementary Fig. 6.**
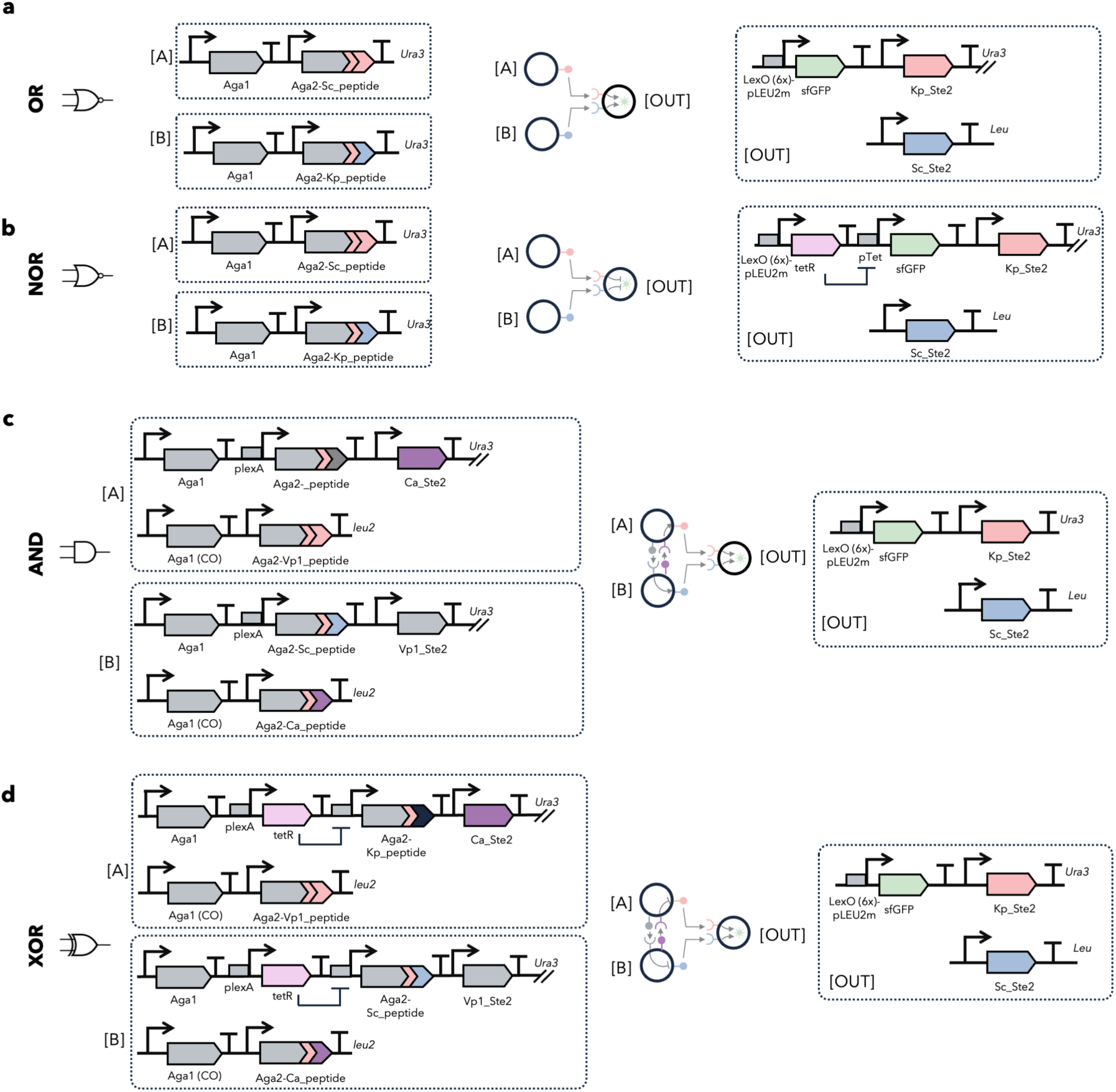
| The genetic circuits designed of four logic gates. The circuits design with key DNA part annotated of four logic OR (a), NOR (b), AND (c) and XOR (d) shown in Figure 2d, 2f, 2h and 2j. Otherwise specified, all the promoters used in in a-d are constitutively expressed promoters described in Table S10.

**Supplementary Fig. 7.**
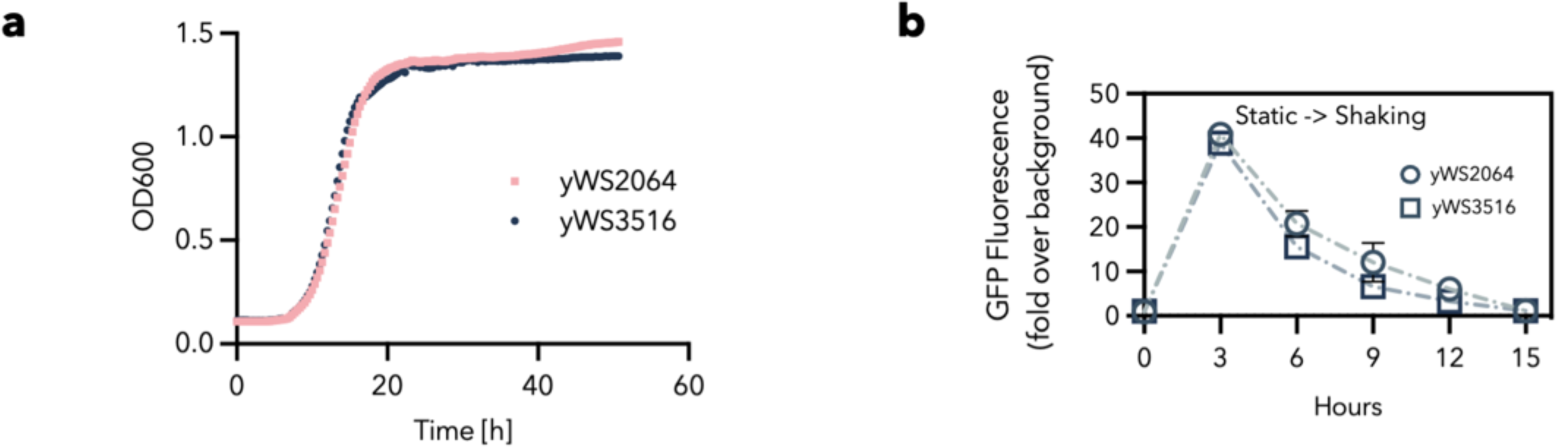
| Characterisation of the base strain, yWS2064 and yWS3516 strains used in this project. a No significant differences in growth rate were observed between yWS2064 and yWS3516 strains when grown in shaking conditions. b In comparison to yWS2064, yWS3516 exhibits a faster return to the resting state of low mean GFP expression when it returns to shaking conditions (700 RPM) after a period in static conditions possibly due to the reduction of native adhesion mediated MARS activation. GFP measurements were conducted in triplicate, and error bars represent the standard deviation.

**Supplementary Fig. 8.**
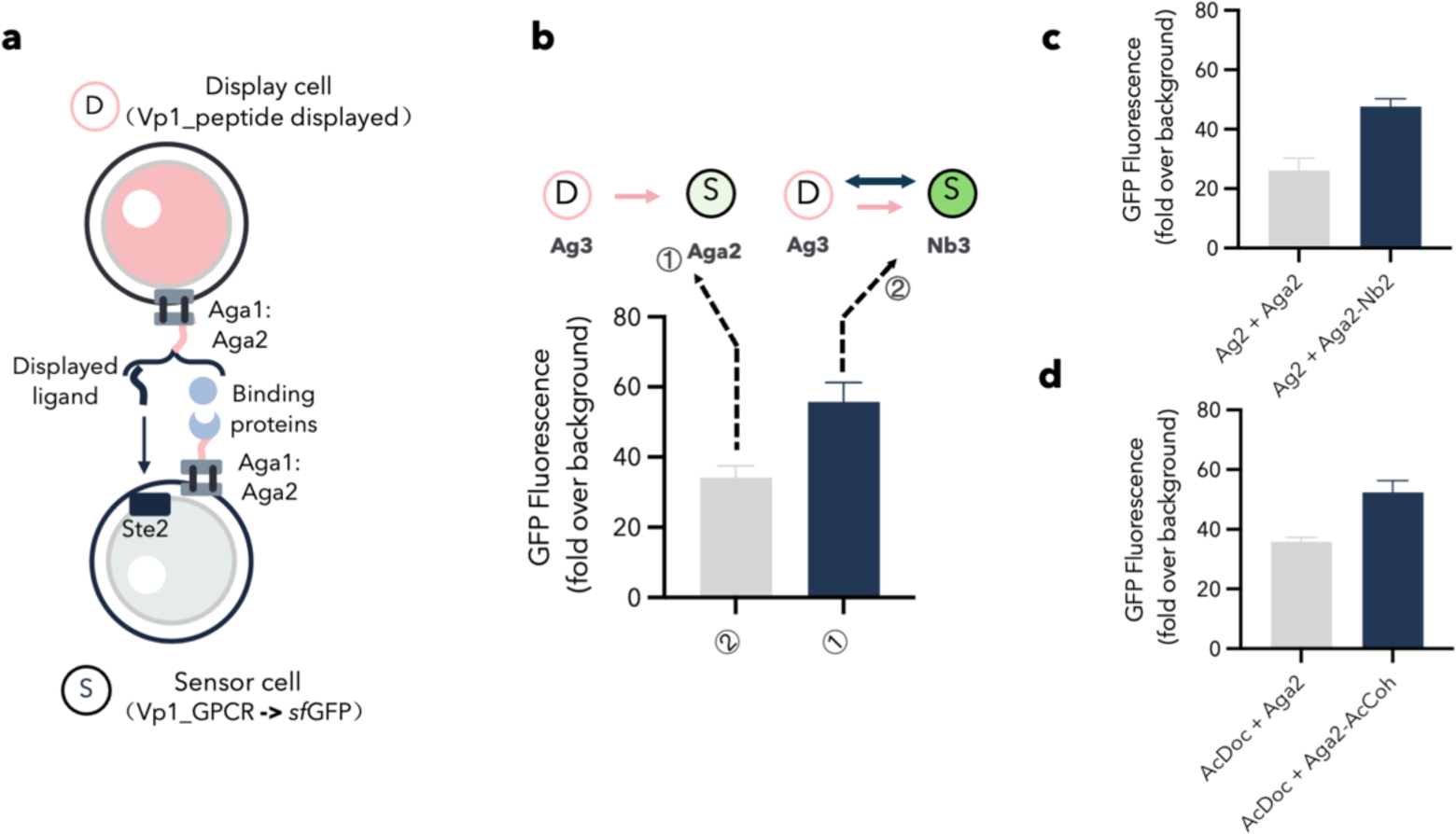
| The effect of adhesive protein pairs on the activation of MARS signalling. a Adhesive protein pairs are displayed on the cell wall using aga1-aga2 surface display, alongside the components used for MARS. b Vp1_MARS with the Ag3-Nb3 adhesion pair added shows a higher sensor cell GFP output level than the control cell with only Ag3-Aga2 displayed. c,d Similar sensor activation profiles were also observed for Vp1_MARS with Ag2-Nb2 displayed (c) and Vp1_MARS with AcDoc-AcCoh displayed (d). Experiments were conducted in triplicate, and all graphs show the mean and error bars representing the standard deviation.

**Supplementary Table 1.**
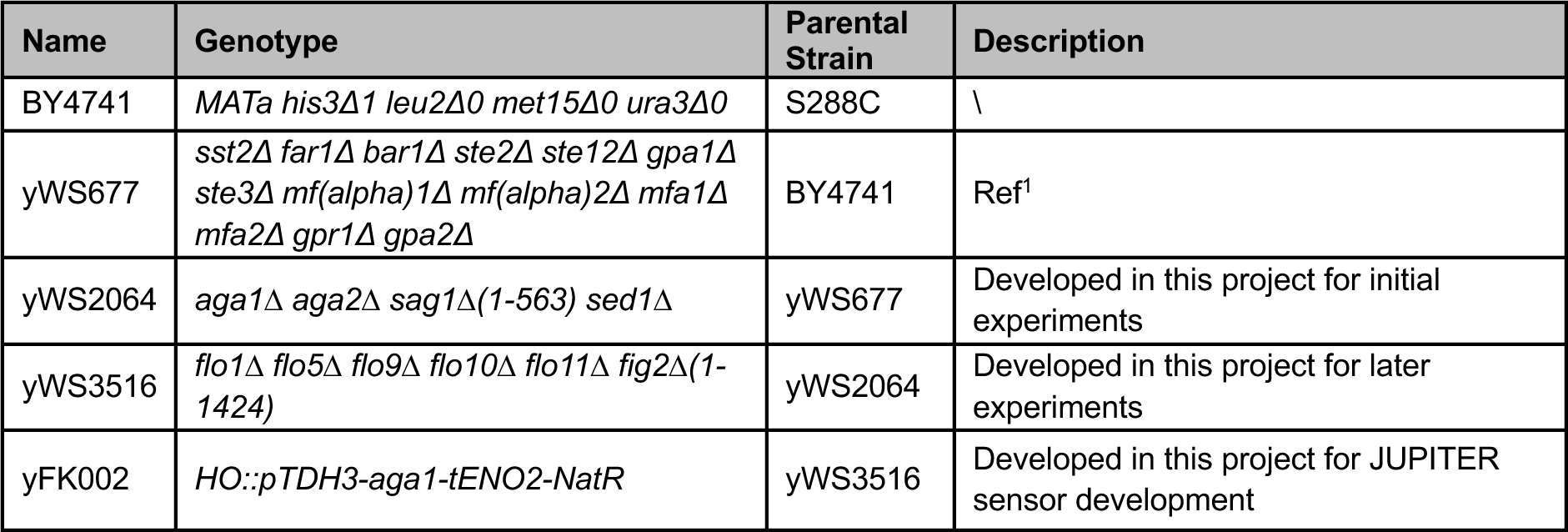
| List of base strains in used in this project.

**Supplementary Table 2.**
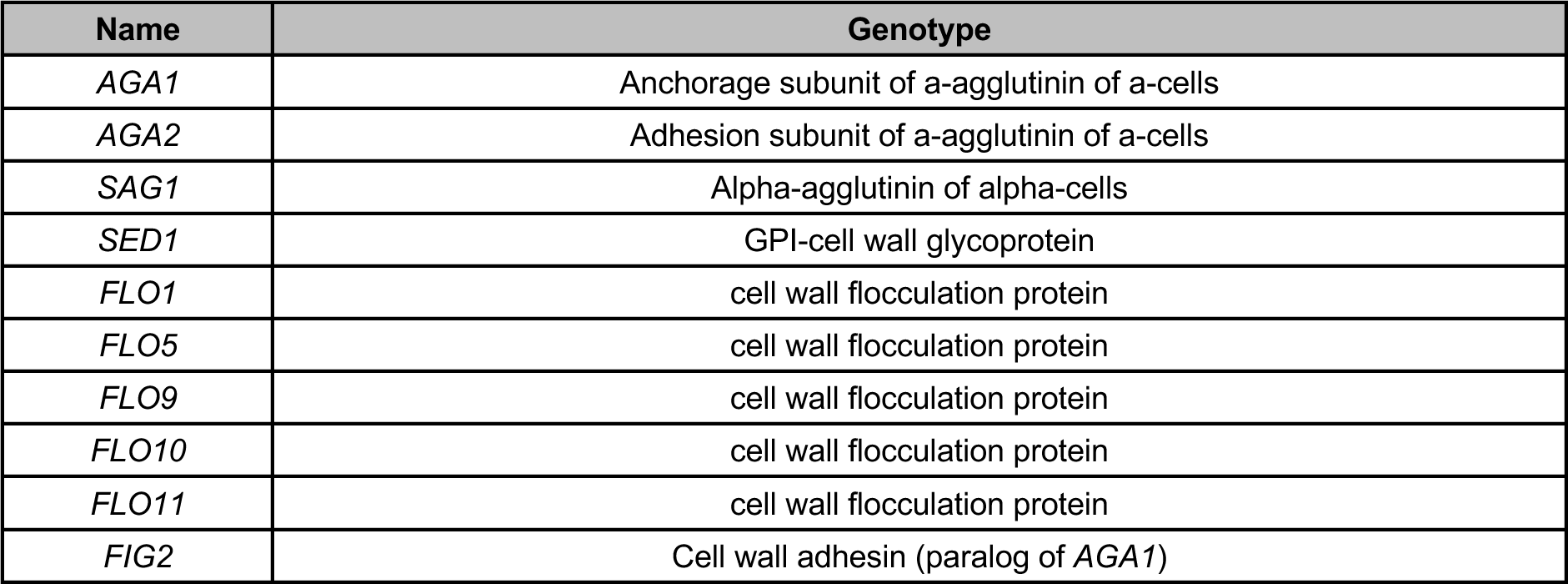
| List of native cell surface display and adhesion related genes deleted to create yWS2064 and yWS3516. Due to the repetitive nature of the sub-telomeric regions surrounding *FLO1*, *FLO5*, and *FLO9*, large deletions were required in yWS3516 to delete these genes, encompassing multiple non-essential, non-functional, or dubious genes (see **Table S3**). No significant differences in growth rate were seen between yWS2064 and yWS3516. *SED1* was deleted to improve the use of the Sed1 protein as an alternative cell surface display anchor but this was eventually not used in the study.

**Supplementary Table 3.**
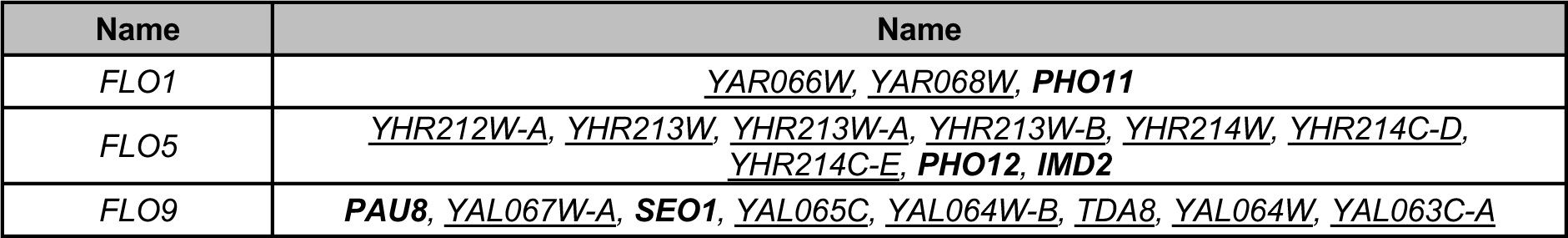
| Genes deleted during the removal of the sub-telomeric regions containing *FLO1*, *FLO5*, and *FLO9*. Dubious ORFs, pseudogenes, and transposable elements are omitted. All verified (bold) and uncharacterised/putative genes are shown.

**Supplementary Table 4.**
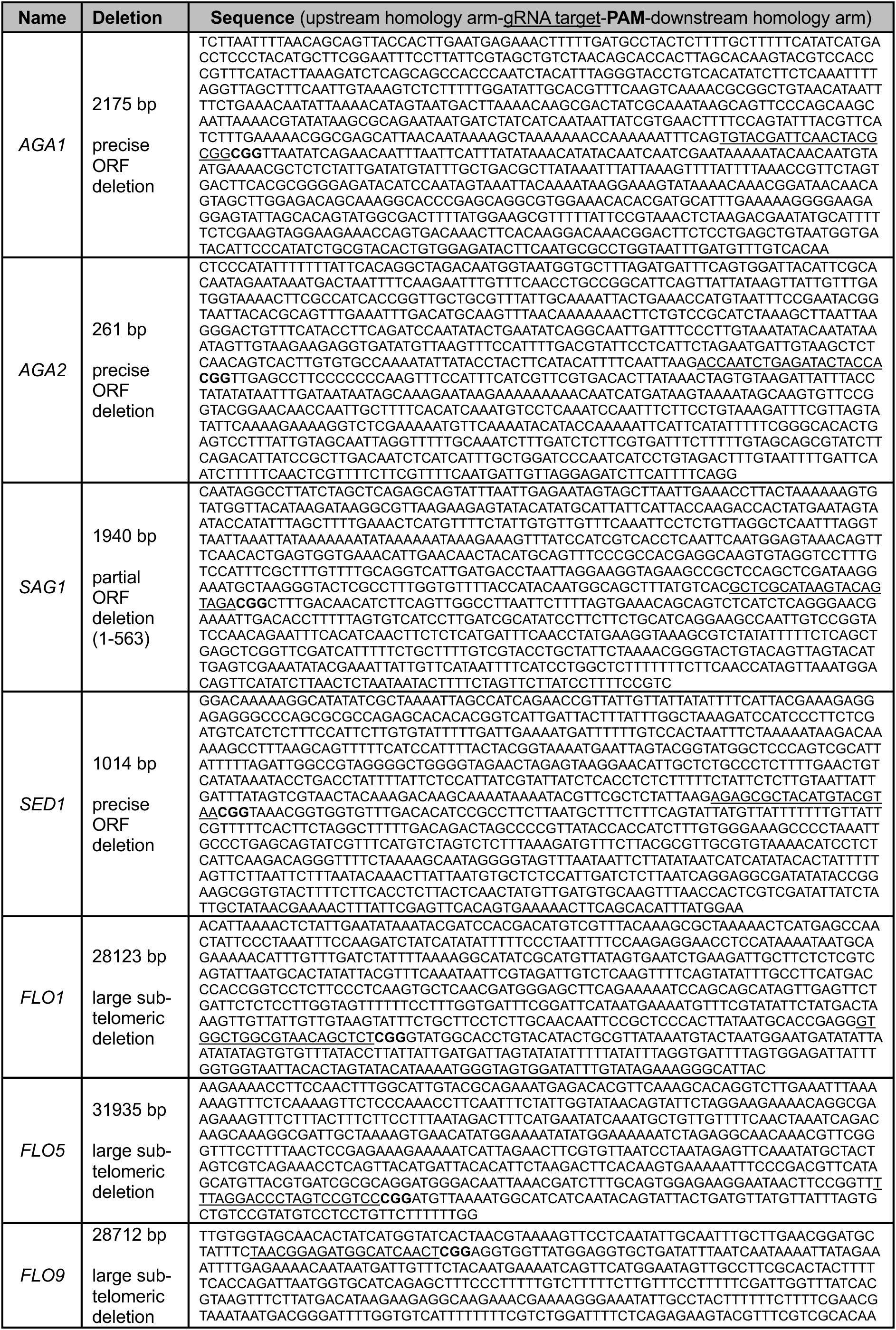
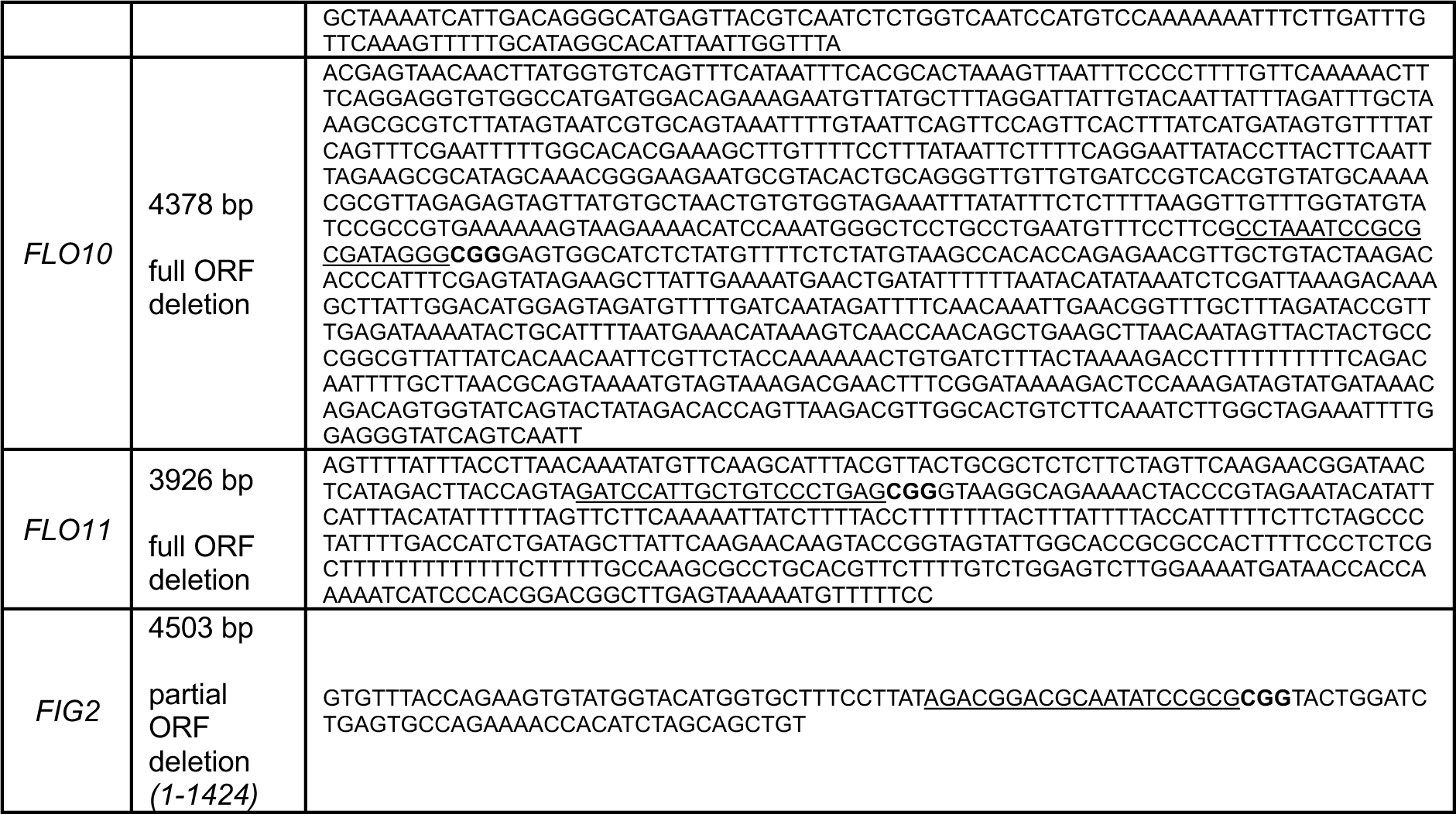
| Donor DNAs used for gene deletions for yWS2064 and yWS3516.

**Supplementary Table 5.**
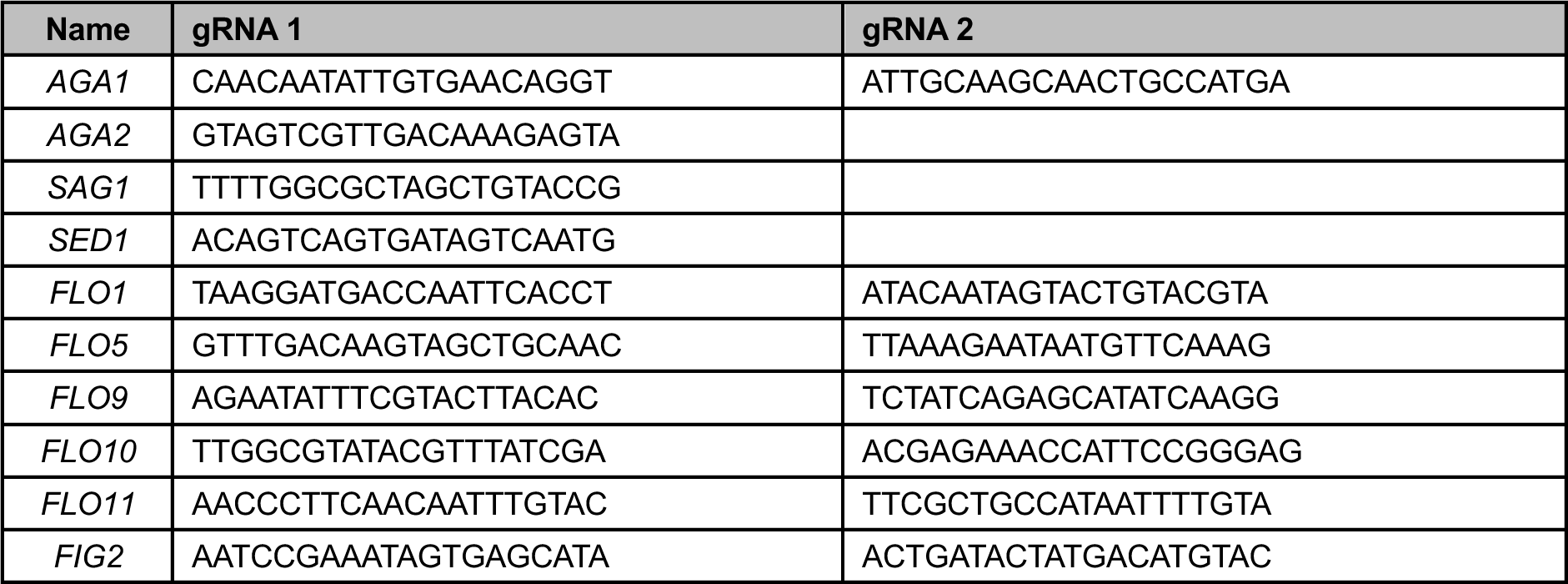
| gRNAs used for gene deletions.

**Supplementary Table 6.**
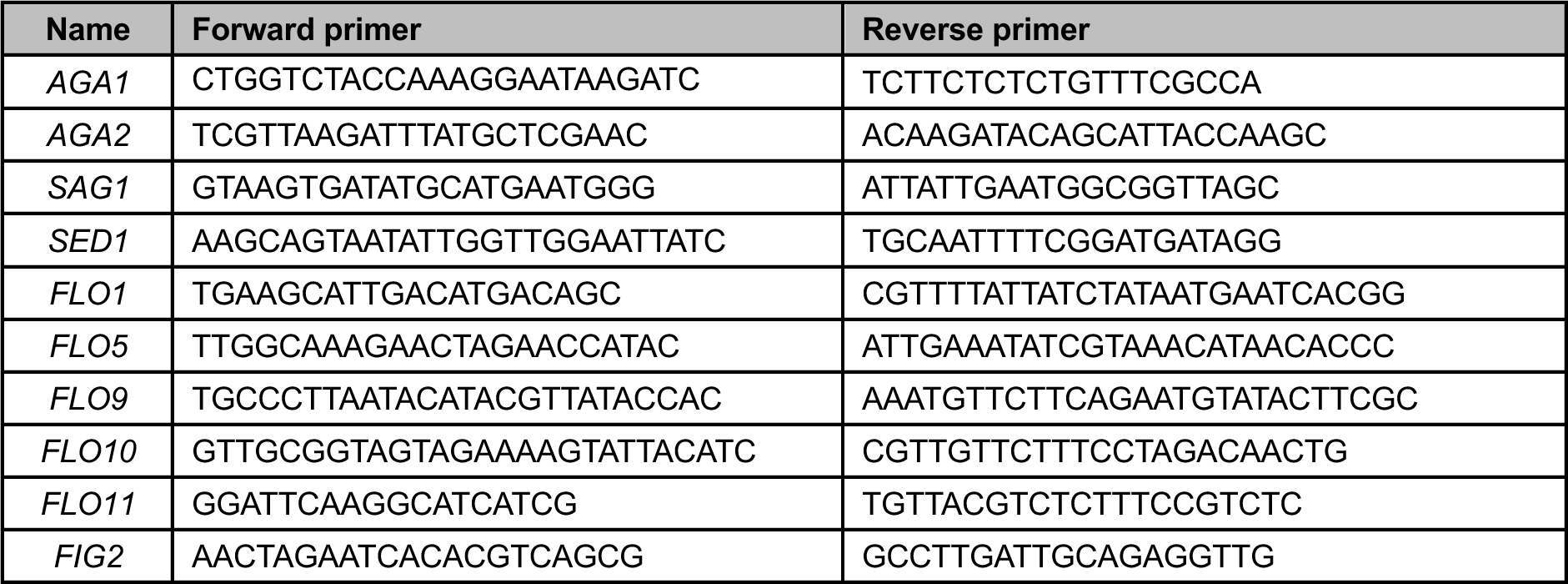
| Gene deletion validation primers.

**Supplementary Table 7.**
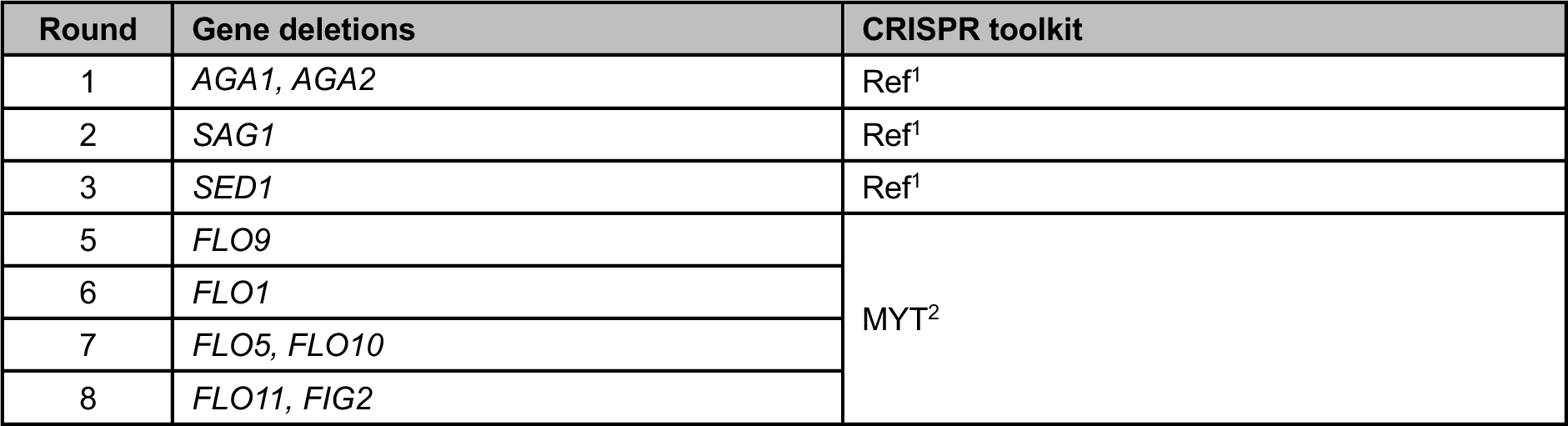
| Gene deletion combinations

**Supplementary Table 8.**
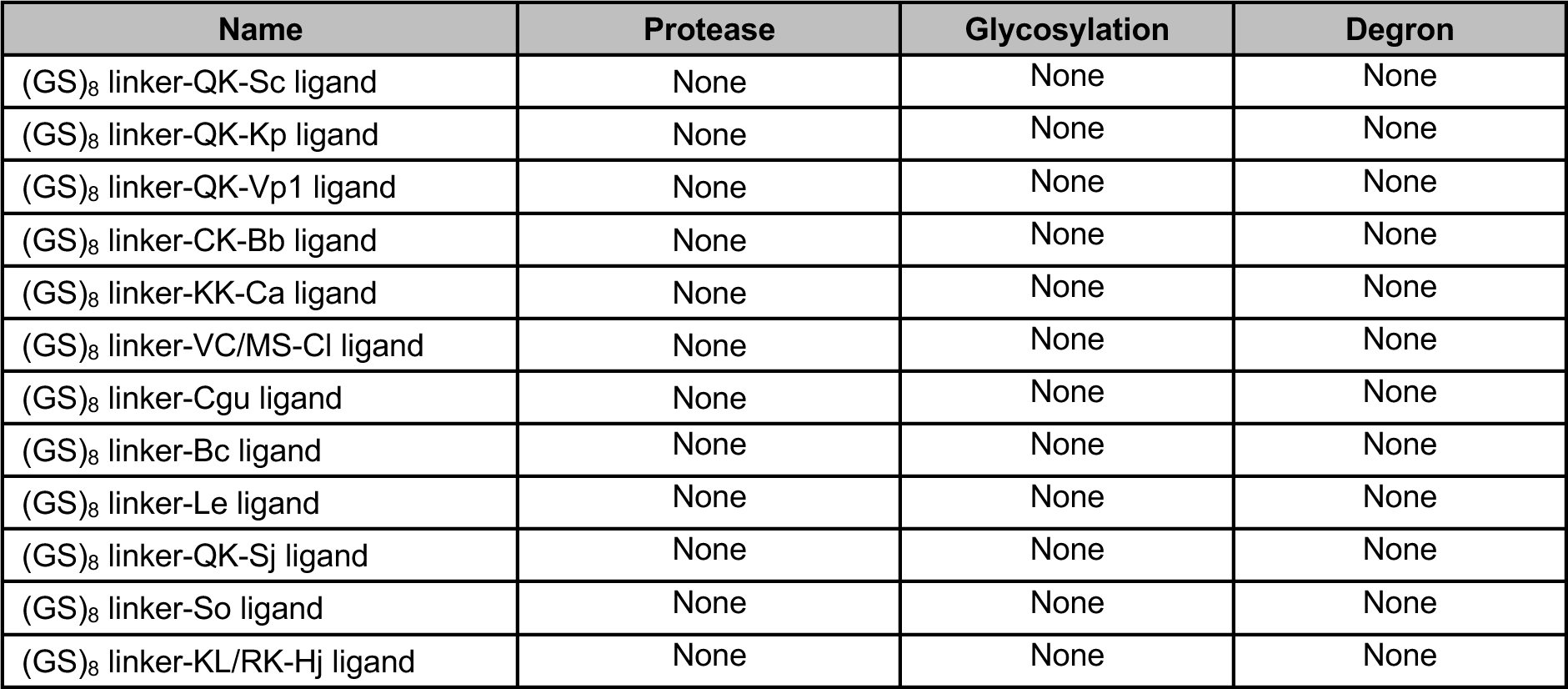
| Eukaryotic Linear Motif (ELM) database results of different ligand display designs

**Supplementary Table 9.**
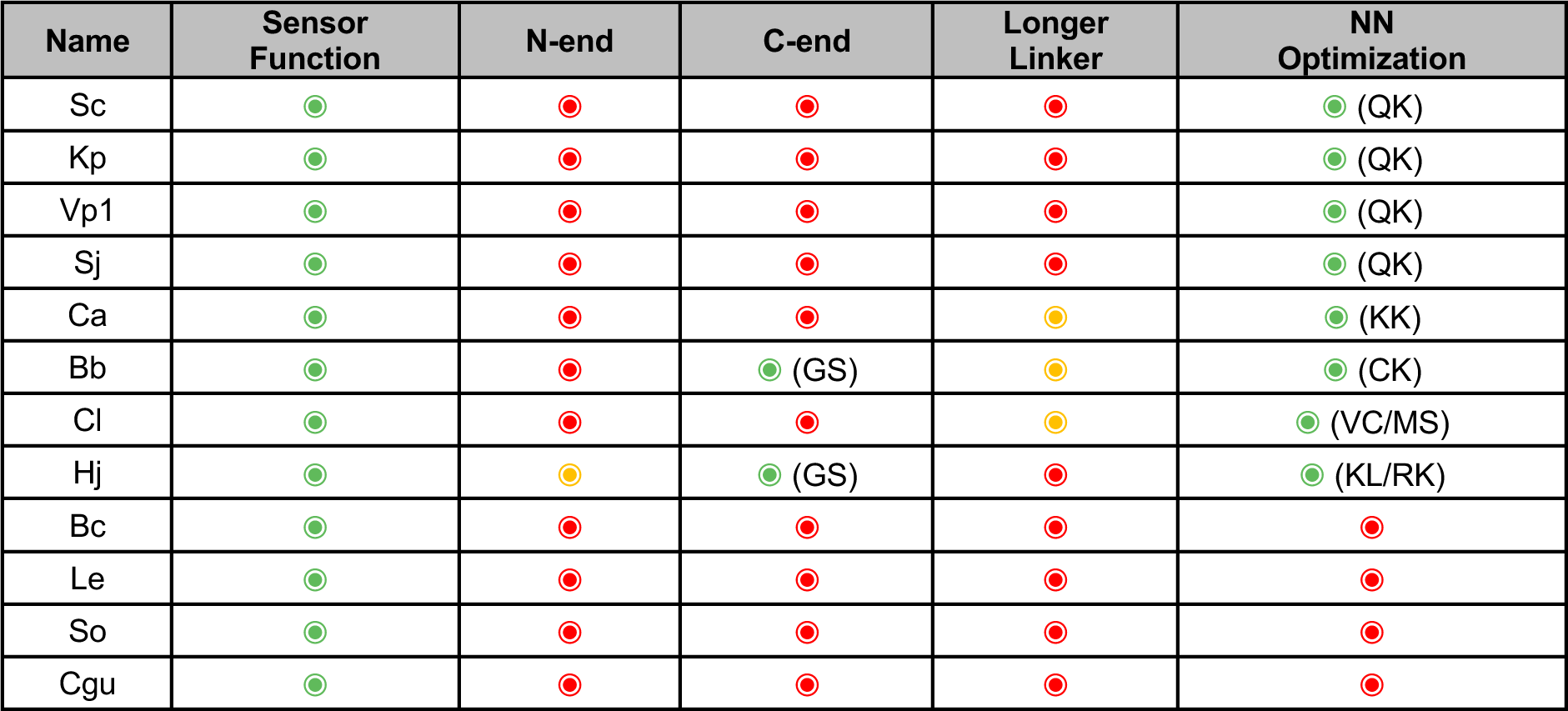
| Overview of different GPCR ligand display strategies

**Supplementary Table 10.**
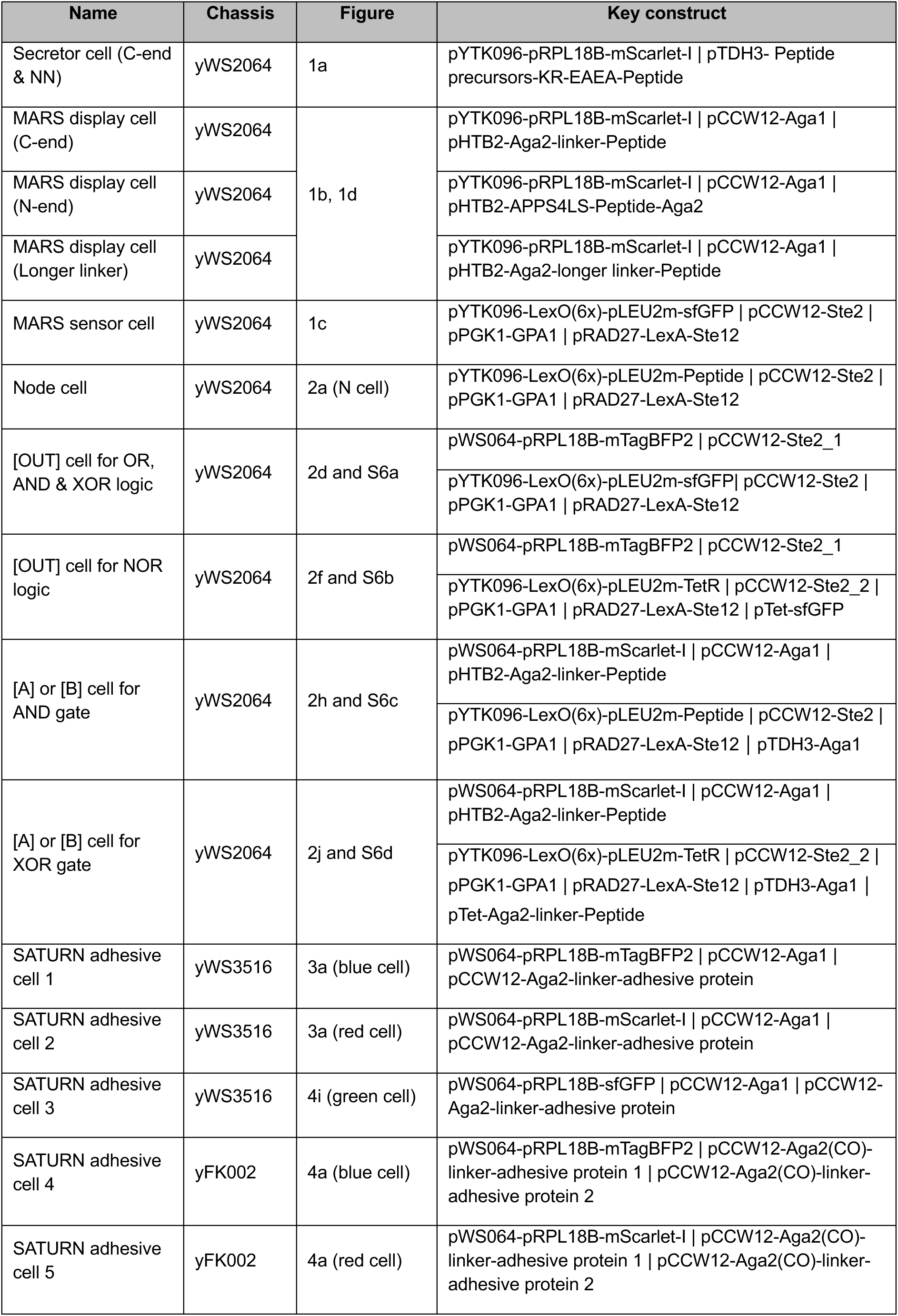
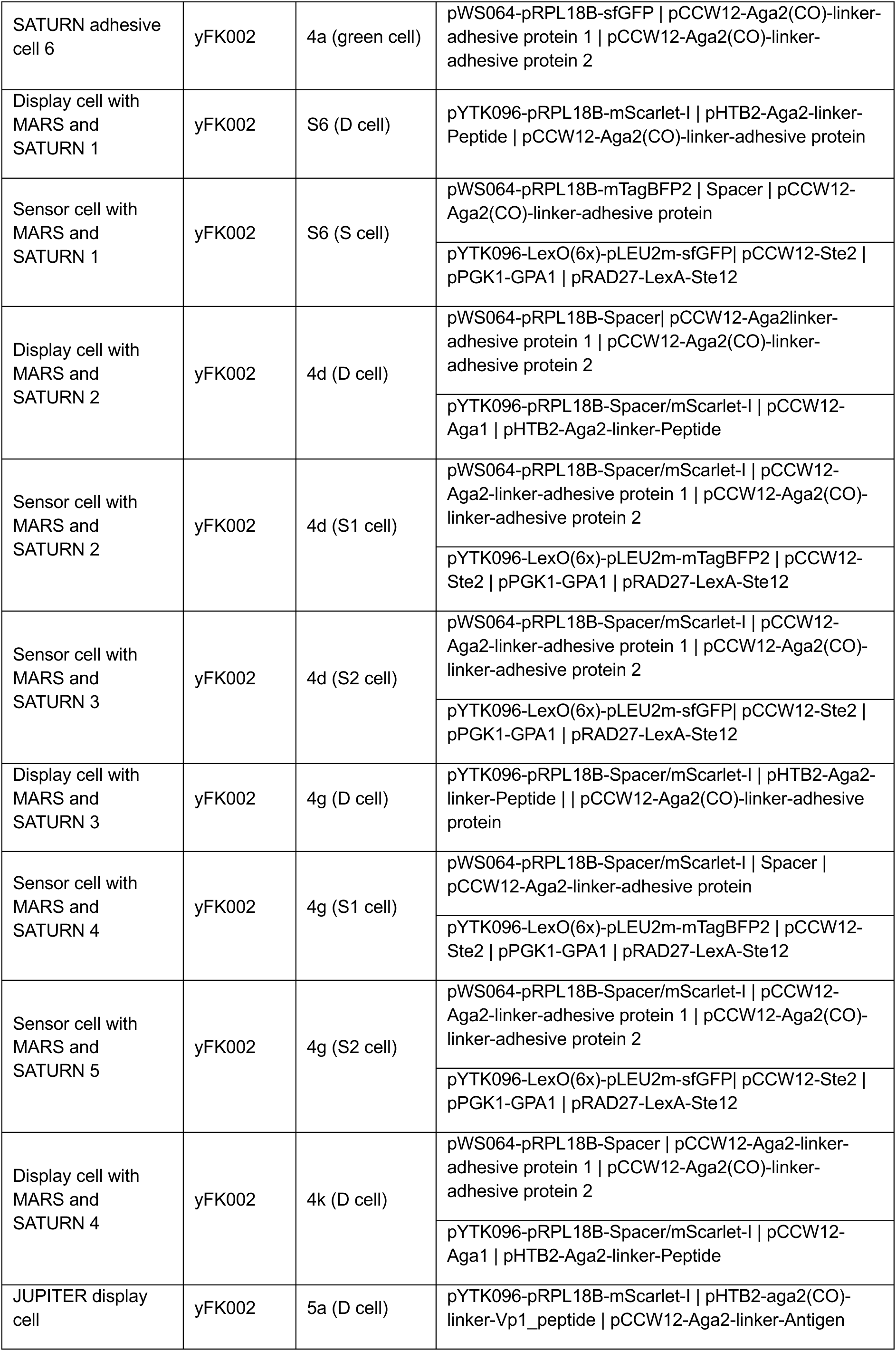
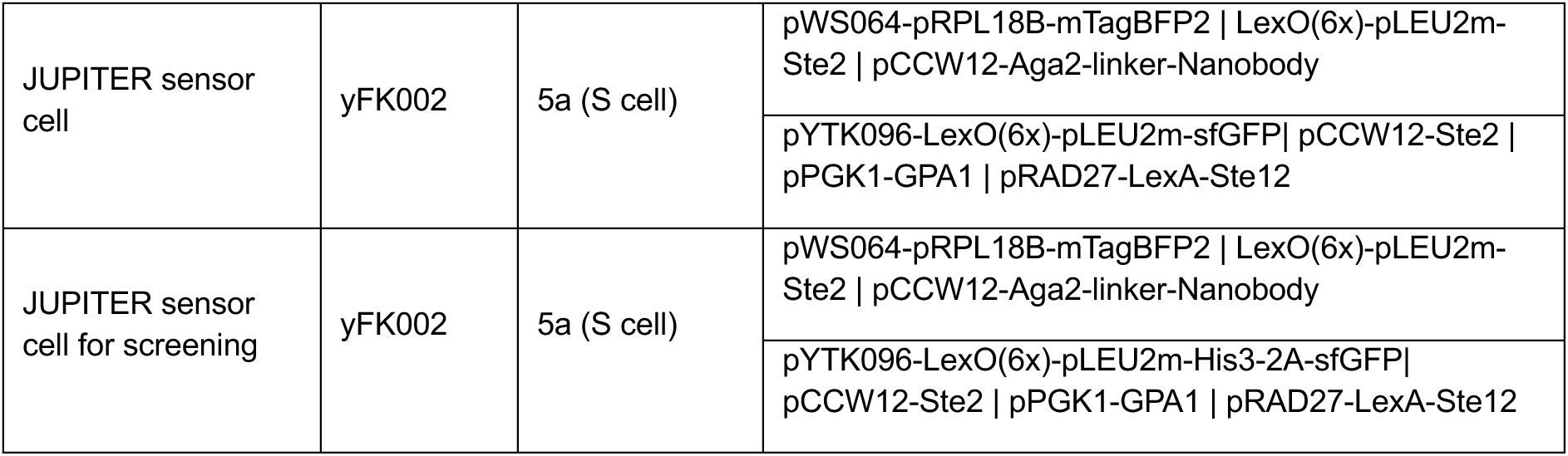
| Key genetic circuits designed in this project.

**Supplementary Table 11.**
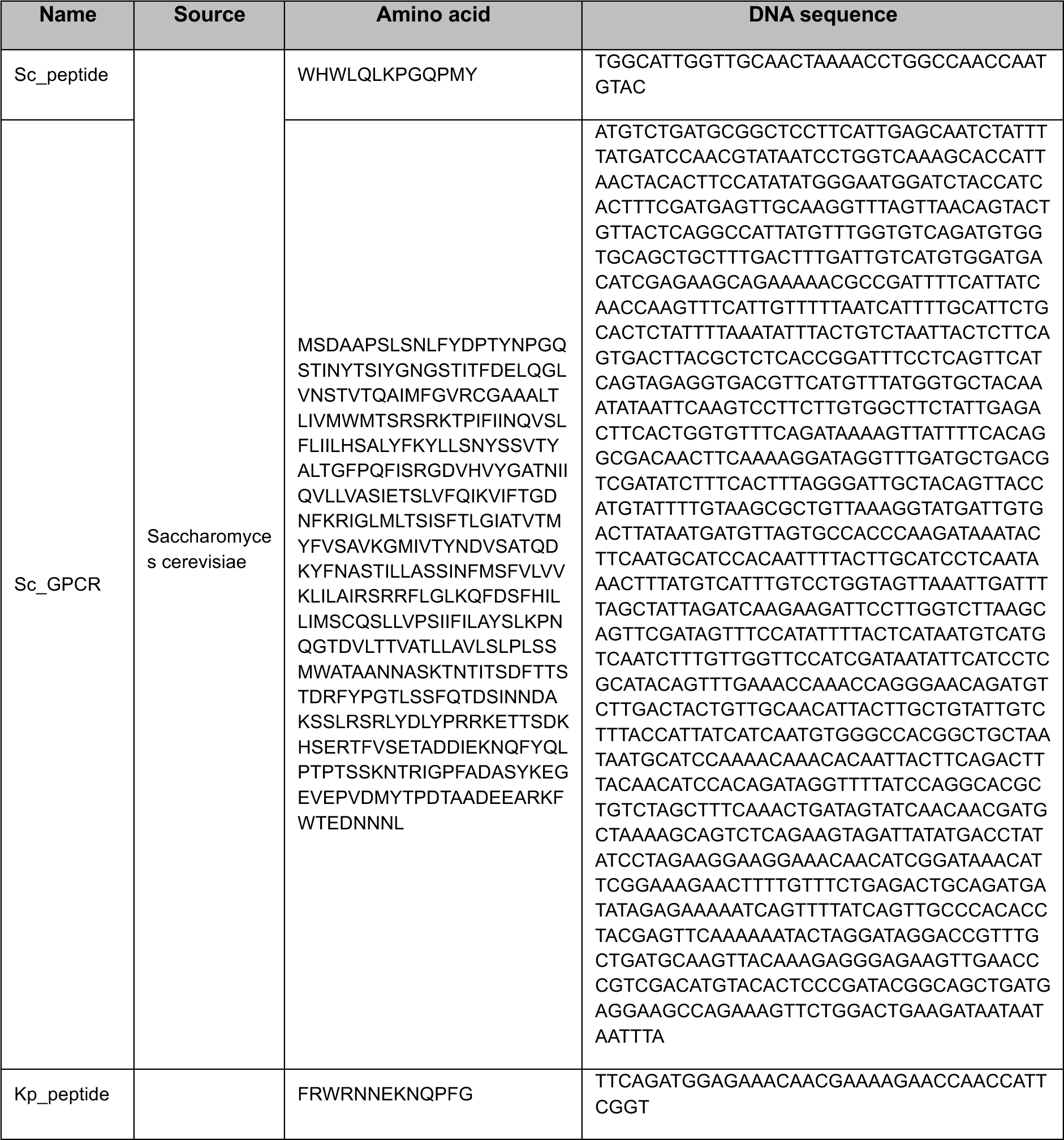
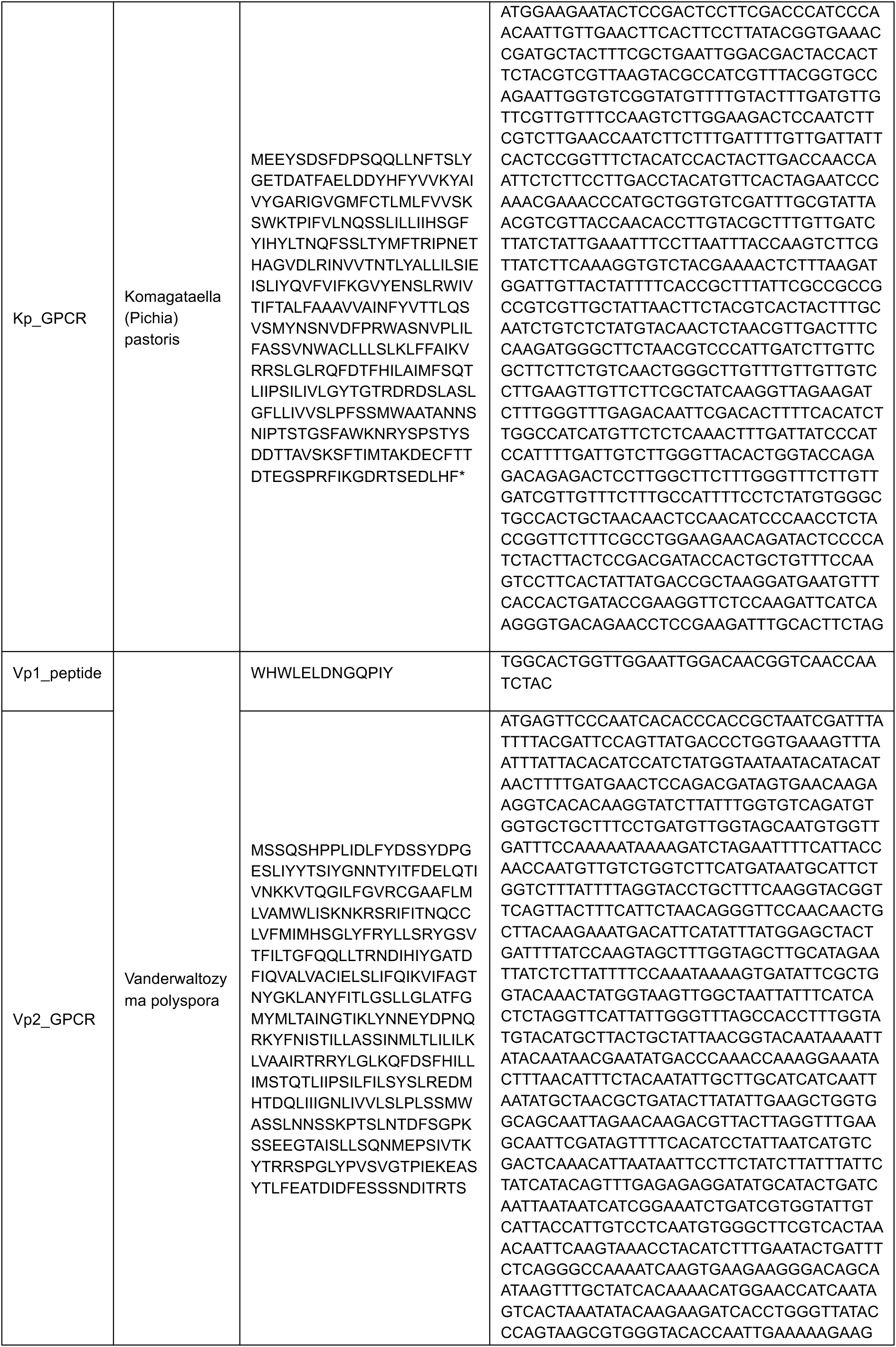
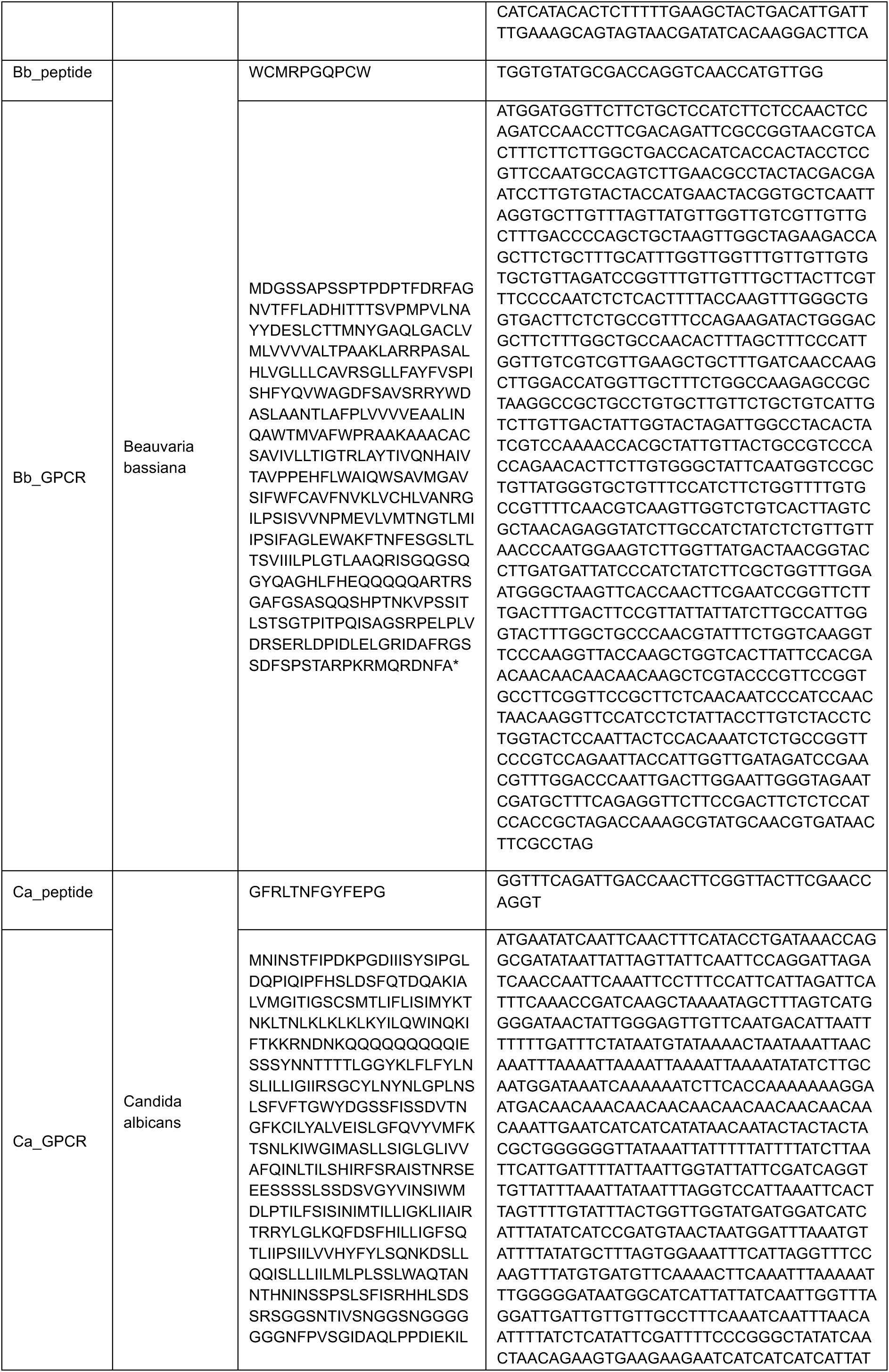
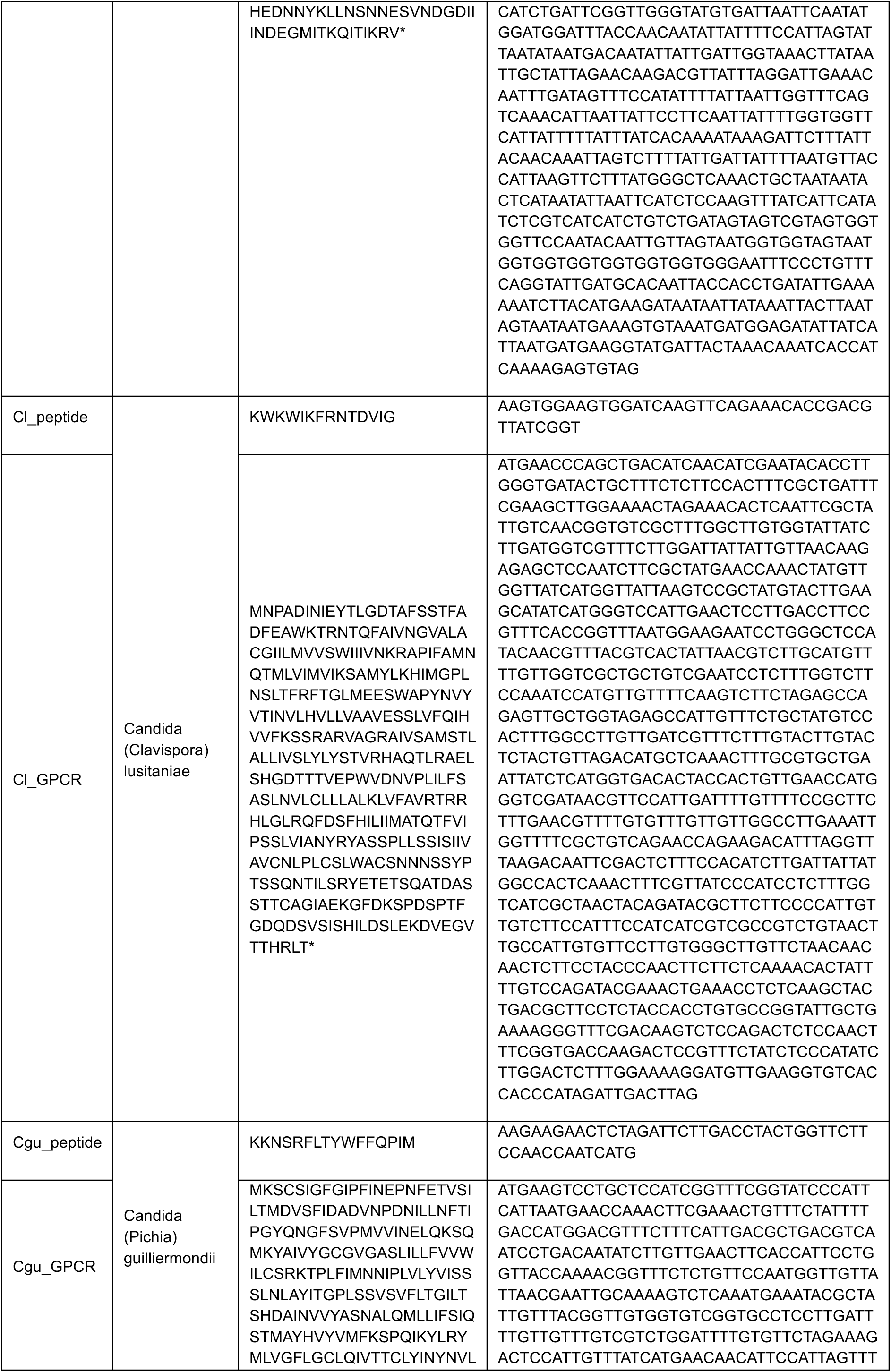
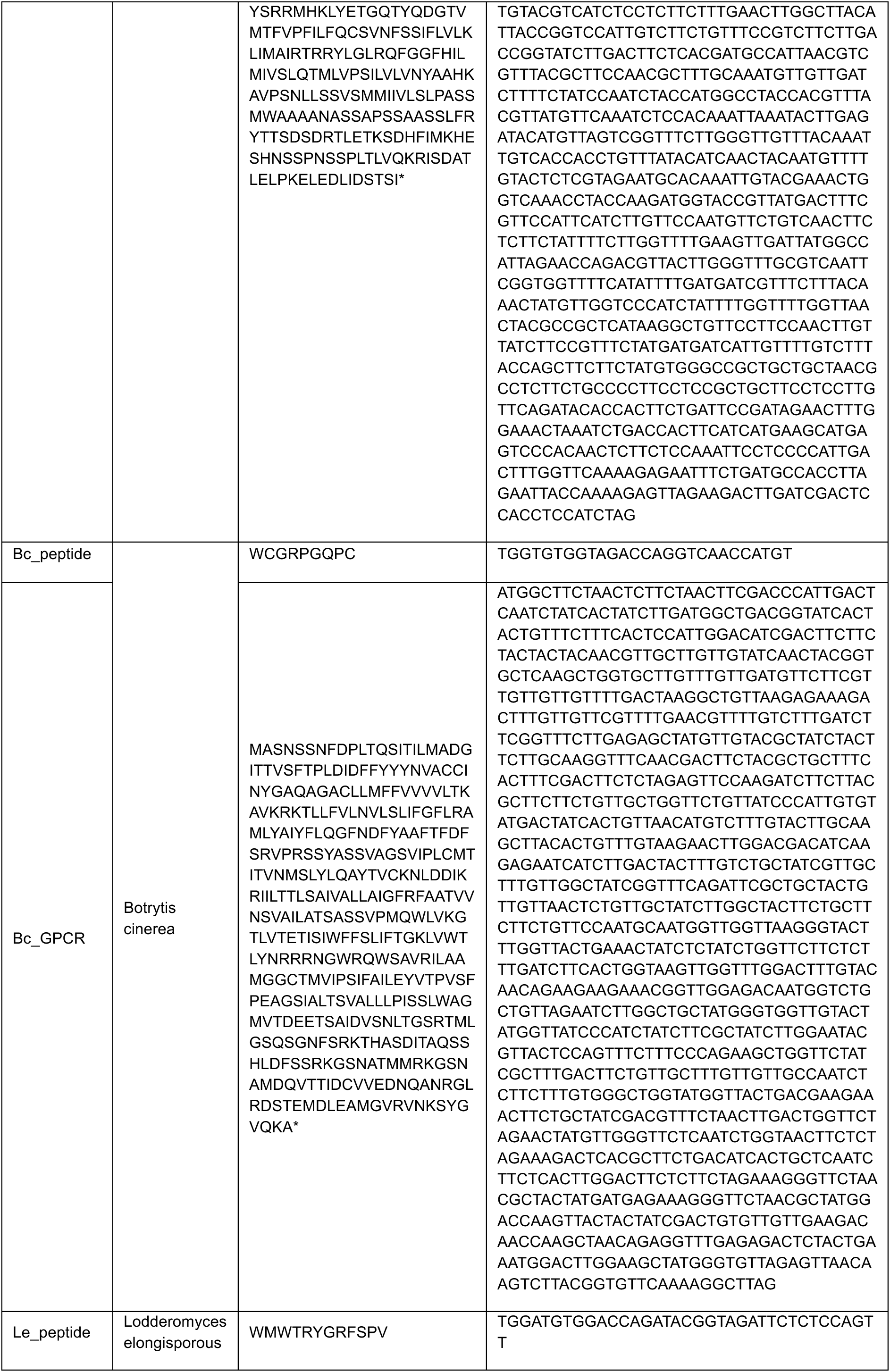
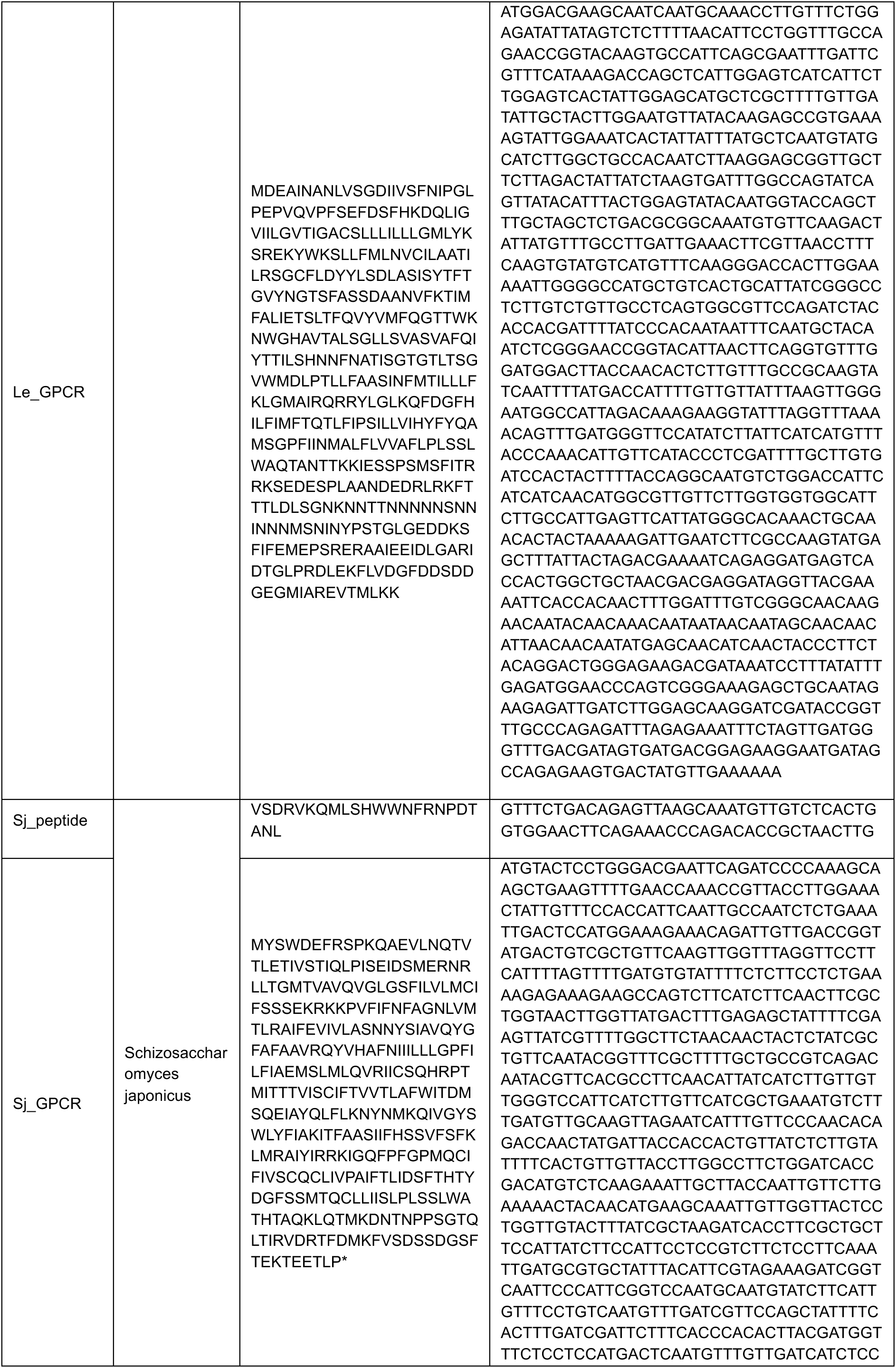
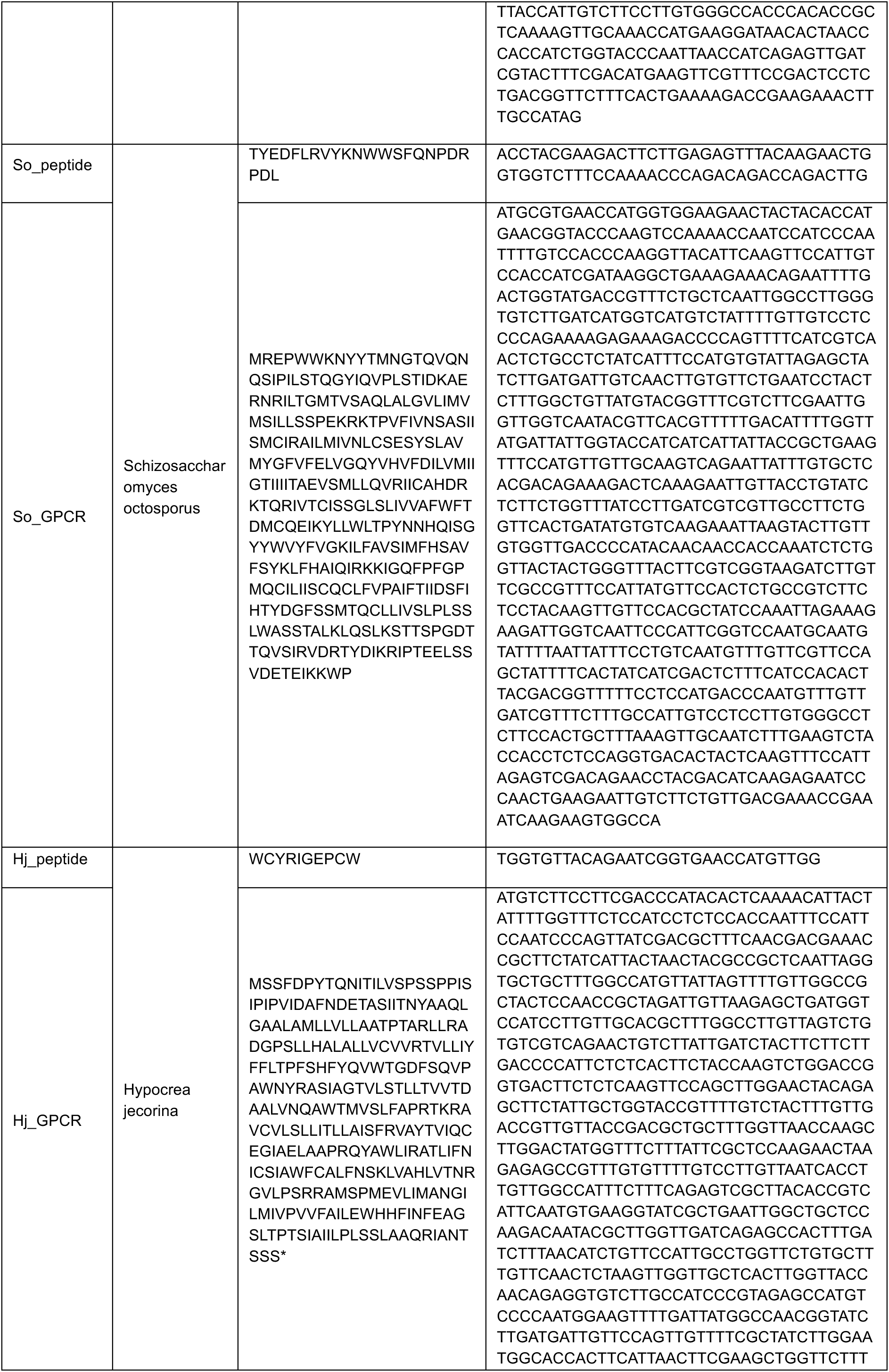
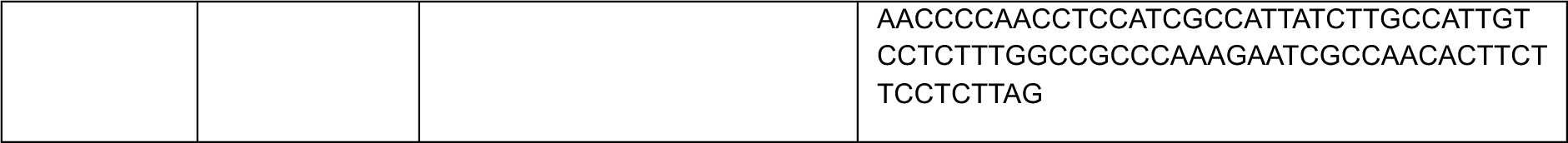
| Sequences of peptide-GPCR ligands used in this project.

**Supplementary Table 12.**
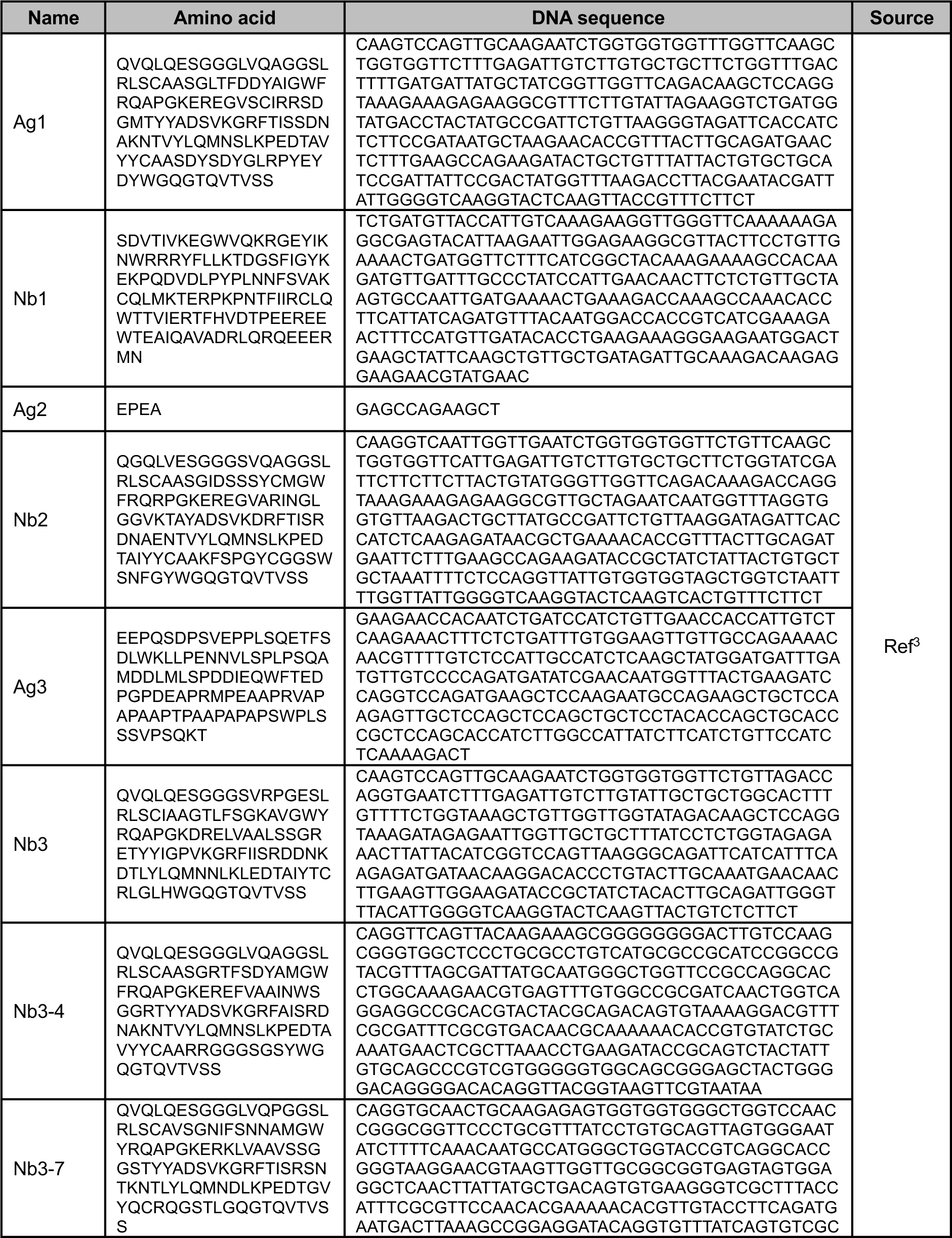
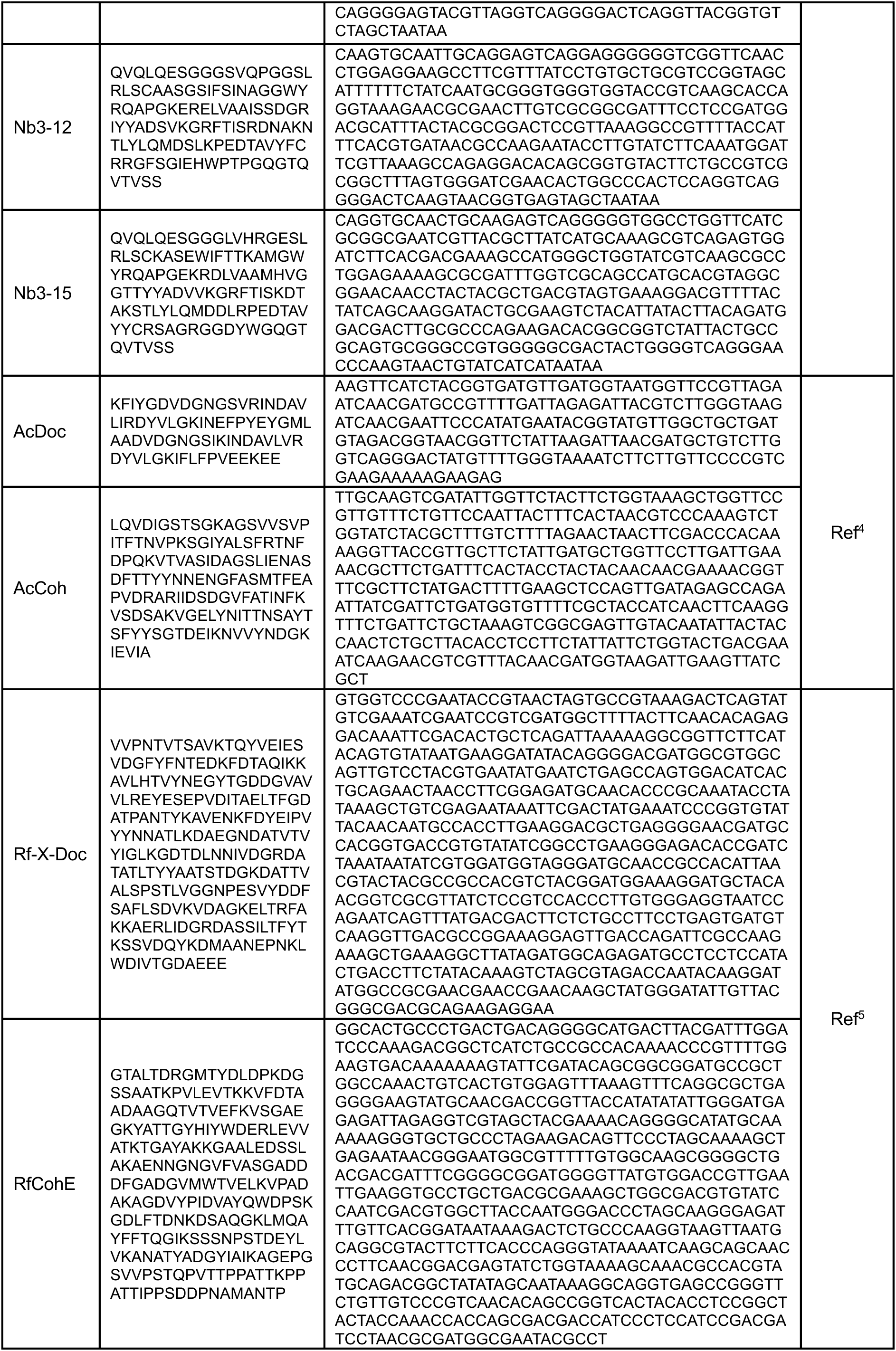
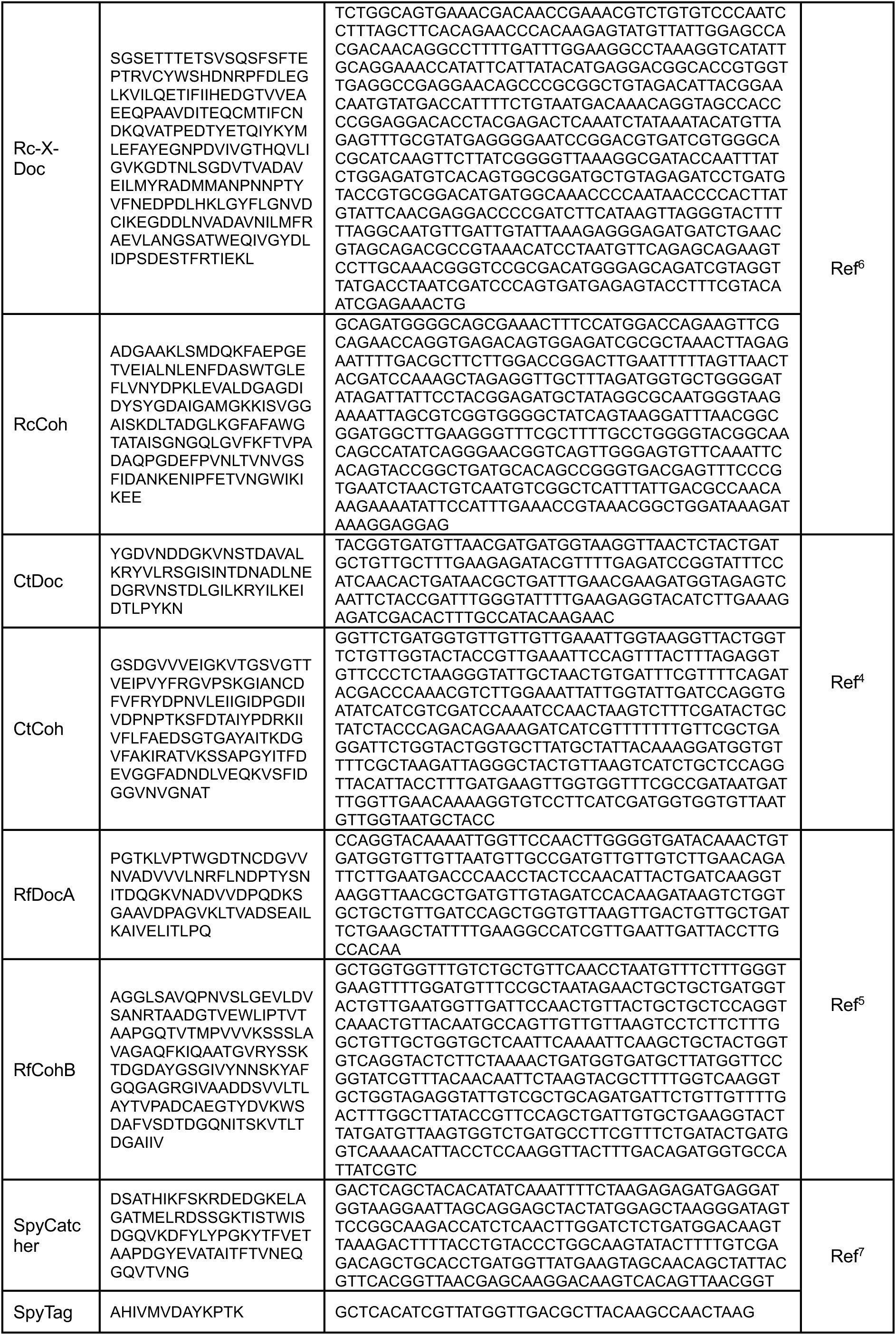
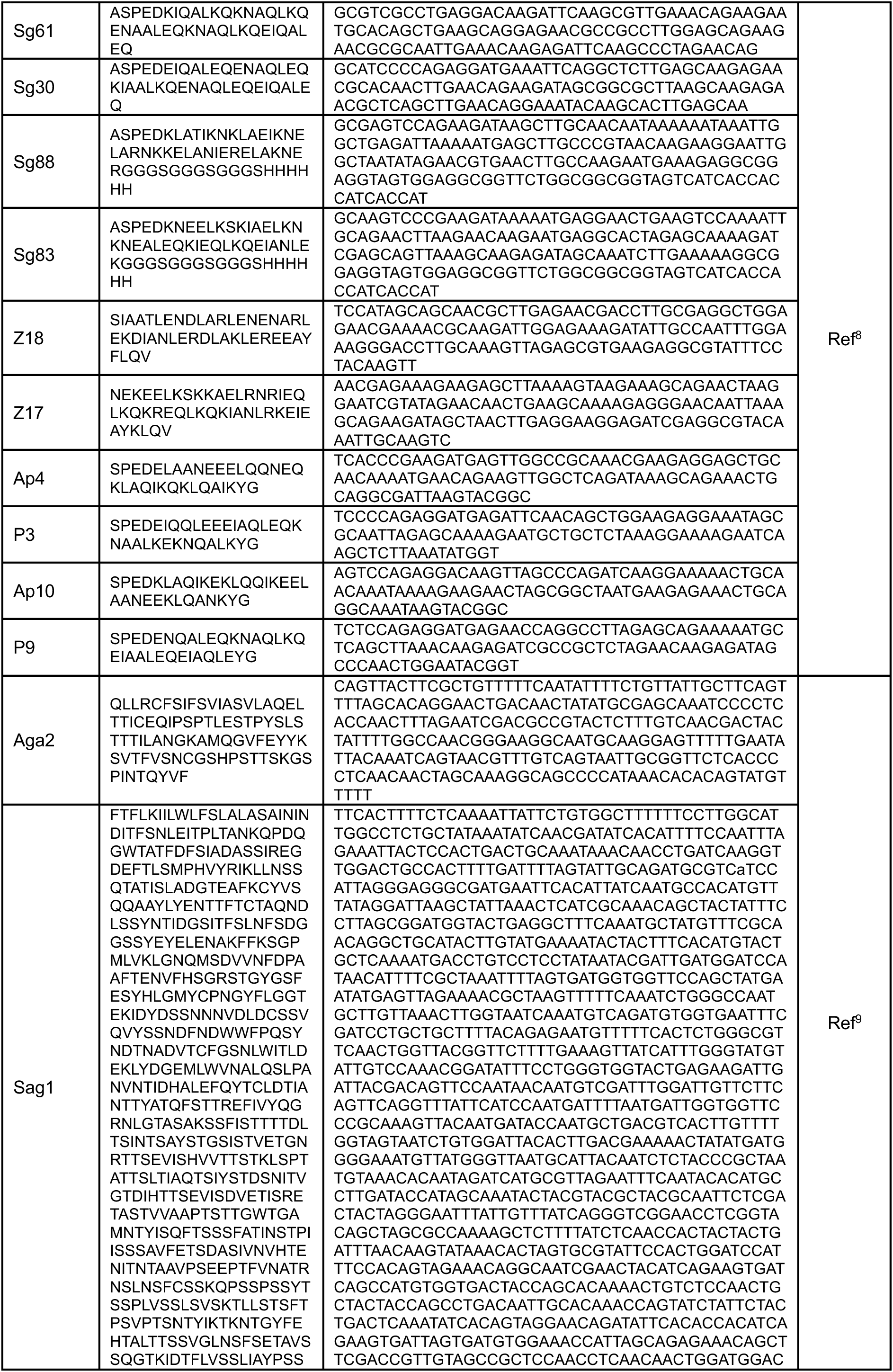
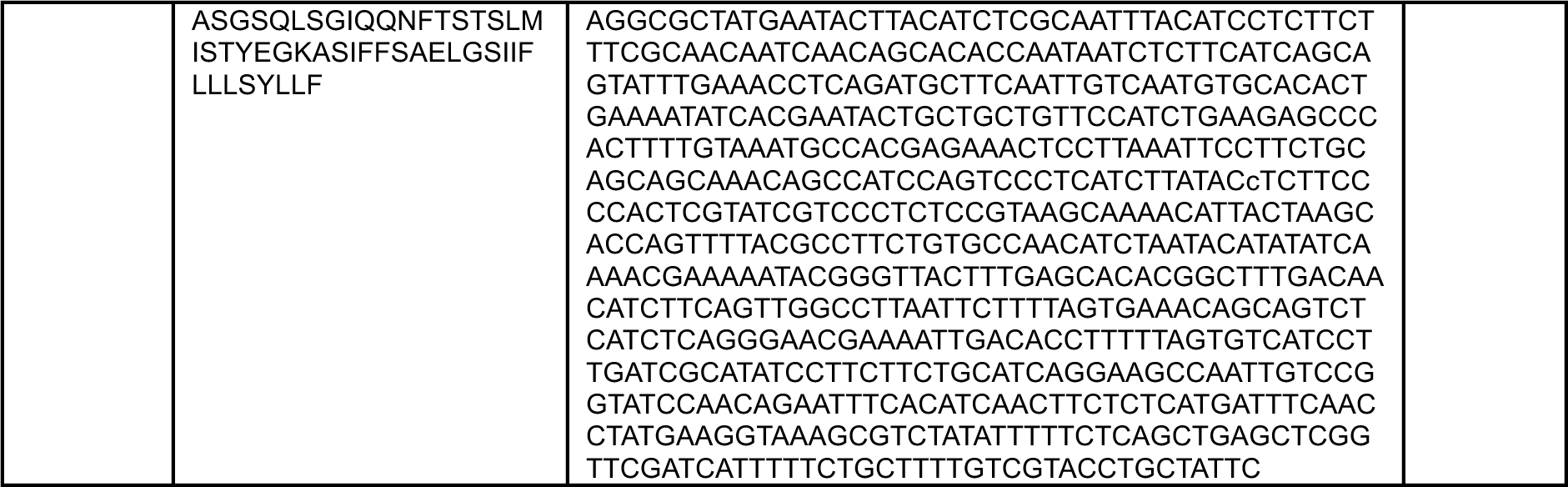
| Sequences of codon-optimized genes for adhesion toolbox development

**Supplementary Table 13.**
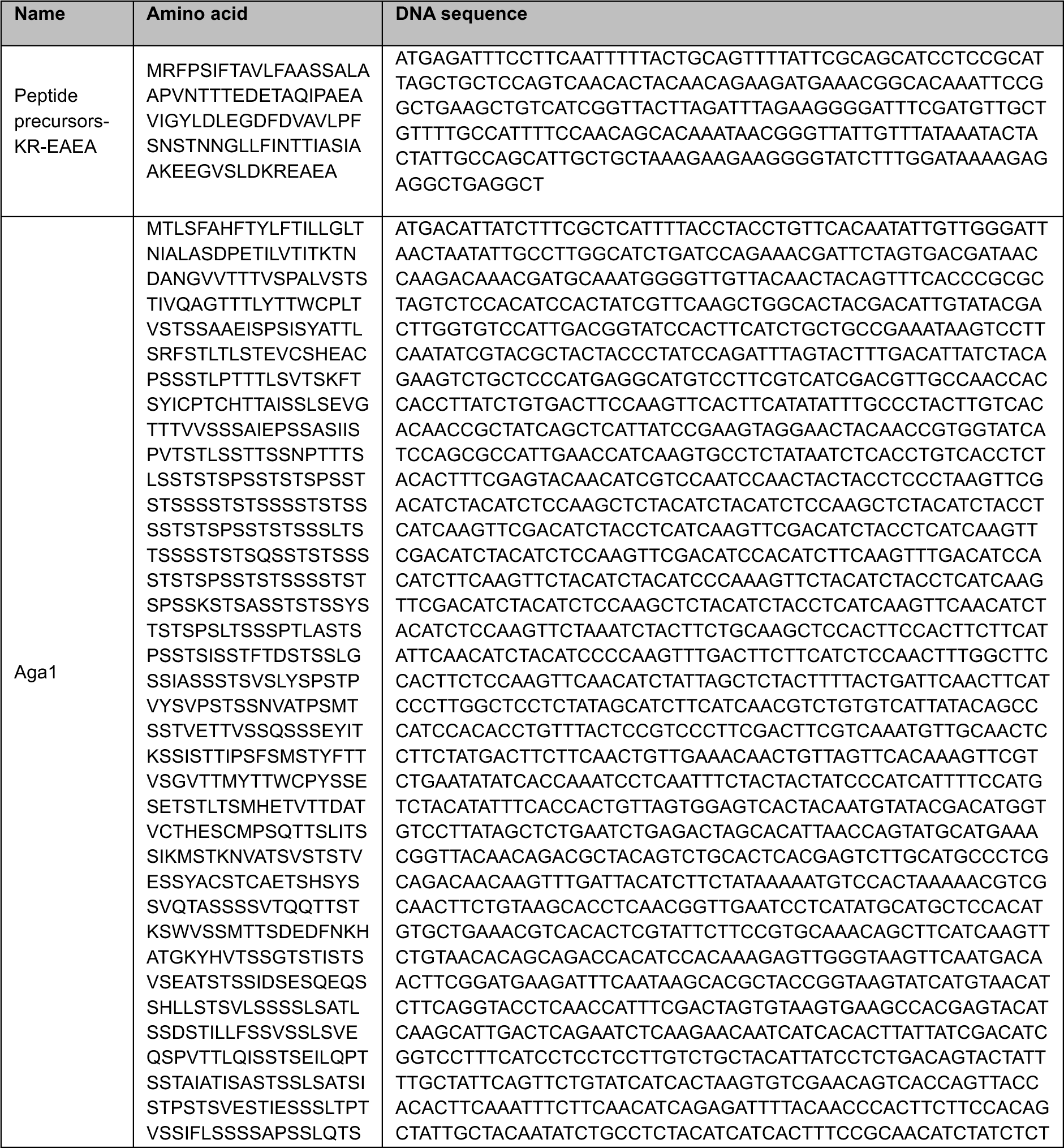
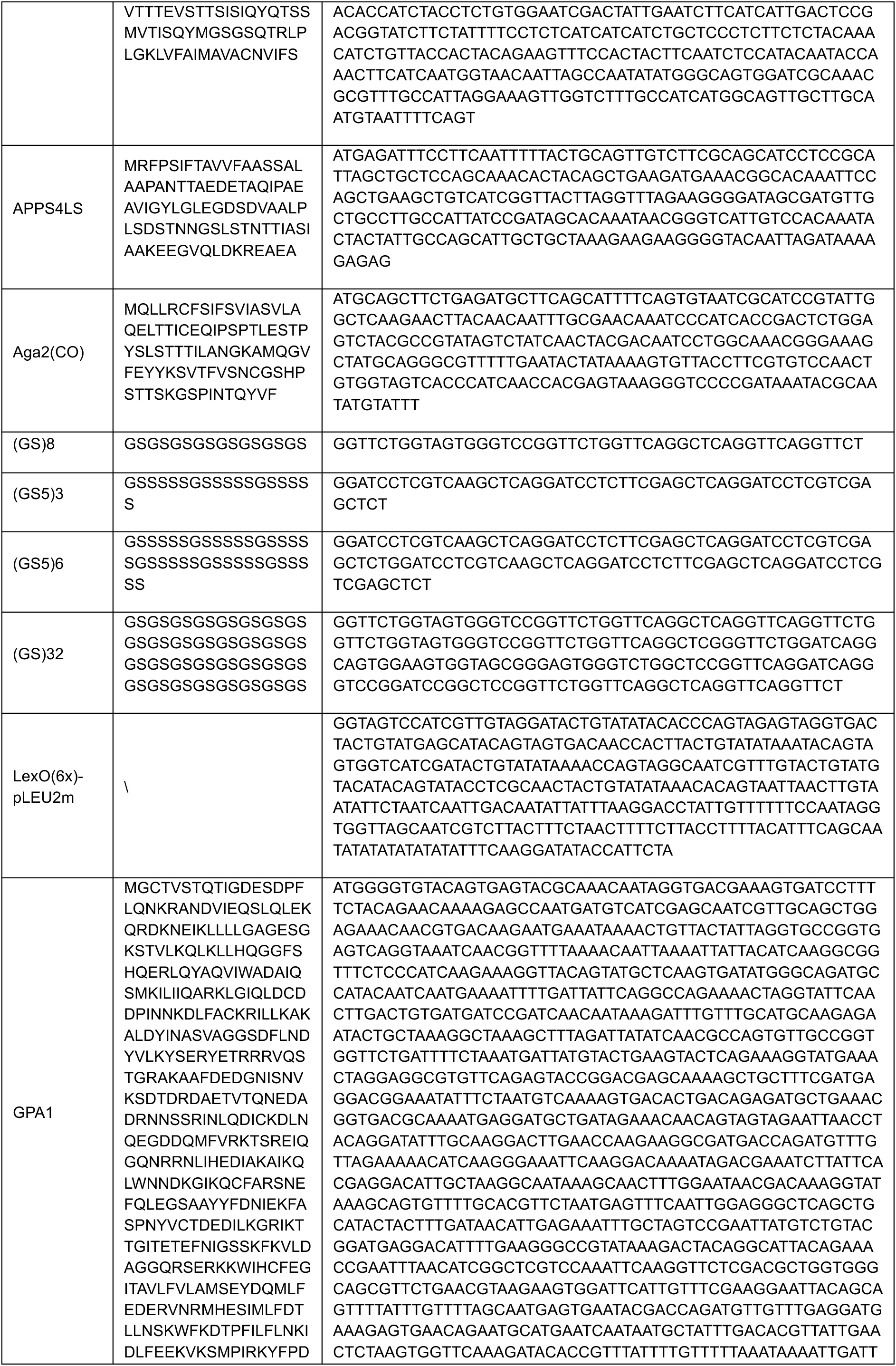
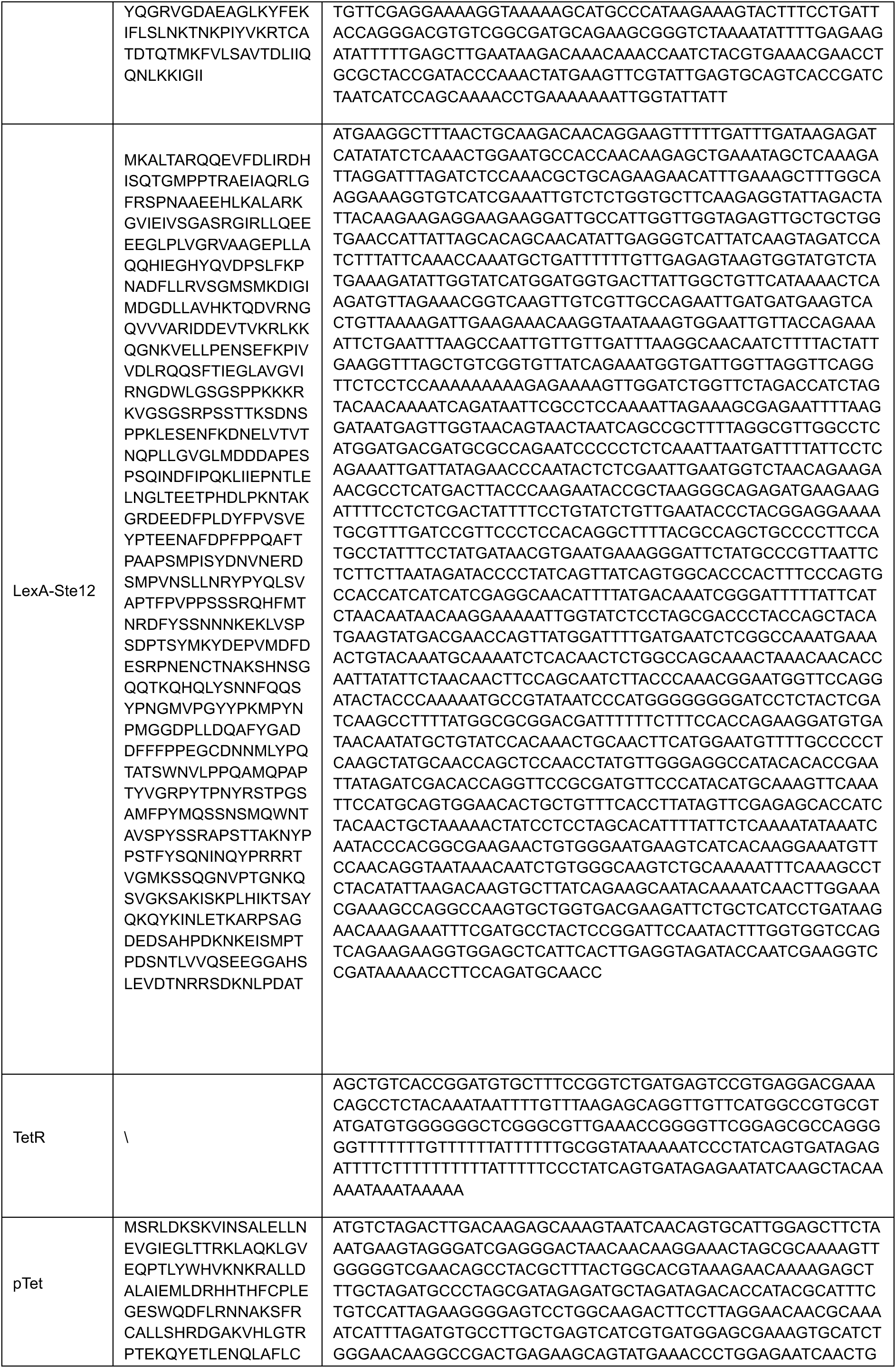
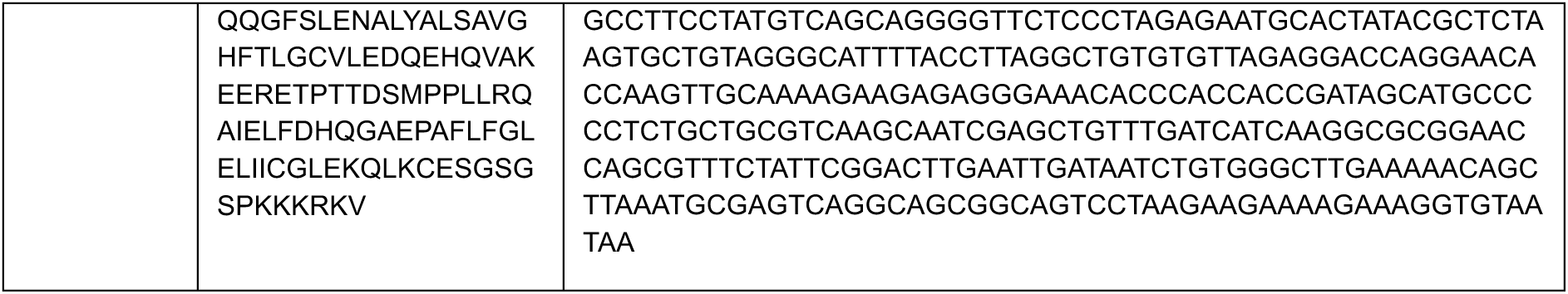
| Summary of key protein and promoter sequence in Table S10.

